# Separable roles of the DNA damage response kinase Mec1^ATR^ and its activator Rad24^RAD17^ within the regulation of meiotic recombination

**DOI:** 10.1101/496182

**Authors:** Margaret R. Crawford, Tim J. Cooper, Marie-Claude Marsolier-Kergoat, Bertrand Llorente, Matthew J. Neale

## Abstract

During meiosis, programmed DNA double-strand breaks (DSBs) are formed by the topoisomerase-like enzyme, Spo11, activating the DNA damage response (DDR) kinase Mec1^ATR^ via the checkpoint clamp loader, Rad24^RAD17^. At single loci, loss of Mec1 and Rad24 activity alters Spo11-dependent DSB formation and recombination outcome, but their genome-wide roles have not been examined in detail. Here we utilise two distinct methods to characterise the roles Mec1 and Rad24 play in meiotic recombination—deletion of the mismatch repair protein, Msh2, and control of meiotic prophase length via regulation of the Ndt80 transcription factor—to enable genome-wide mapping of meiotic progeny. In line with previous studies, we observe a reduction in the frequency of recombination upon deletion of *RAD24*—driven by a shortened prophase. By contrast, loss of Mec1 function increases recombination frequency, consistent with its role in DSB *trans*-interference. Despite this difference in recombination rate, complex, multi-chromatid events initiated by closely spaced DSBs—a rare event in wild type cells—occur more frequently in the absence of either Rad24 or Mec1, suggesting a loss of spatial regulation at the level of DSB formation. We further demonstrate that Mec1 and Rad24 also have important, yet distinct, roles in the spatial regulation of crossovers (COs). Specifically, excess DSBs forming in the absence of Mec1 are disproportionately channelled into the non-crossover (NCO) and non-interfering CO pathways, reducing the global strength of interference without an explicit change in the frequency of interfering COs. In direct contrast, loss of Rad24 weakens interference via a reduction in the apparent number of interfering COs—similar, but less extreme, to the phenotype of ‘ZMM’ mutants such as *zip3*Δ. Collectively, our results highlight novel and unique roles for Rad24 within meiotic recombination—beyond those mediated by activation of Mec1—and describe new roles for the DDR in several important aspects of meiosis.

## Introduction

Meiosis is a specialized form of cell division that produces haploid cells for sexual reproduction. Integral to meiosis is the process of genetic recombination, which is initiated by programmed DNA double-strand breaks catalysed by the topoisomerase II-like enzyme, Spo11 [1]. Meiotic recombination is monitored by the DNA damage response (DDR) in a similar manner to DNA lesions arising within vegetative cells, likely due to the potentially dangerous nature of DSBs. In the generalised pathway, DNA damage leads to the activation of Tel1^ATM^ (Ataxia Telengiectasia-Mutated) and Mec1^ATR^ (Rad3-related) checkpoint kinases (Human orthologues are indicated with superscript text), via the Mre11-Rad50-Xrs2^NBS1^-complex and Rad24^RAD17^ (reviewed in [2]). Rad24^RAD17^ is the loader of the Ddc1^RAD9^-Rad17^RAD1^-Mec3^HUS1^ (“9-1-1”) checkpoint clamp, that binds to the ssDNA/dsDNA junctions that arise following resection of DNA ends [3]. Tel1^ATM^ and Mec1^ATR^ modulate downstream targets via Rad53^CHK2^ [4], causing cell cycle arrest and the modulation of transcription [5]. Notably, in mammalian cells, RAD17 and ATR have both overlapping and distinct functions in the DDR. RAD17 is required for the recruitment of RAD9 after DNA damage, a function that it performs even in the absence of ATR [6]. Conversely, ATR can localize to sites of DNA damage independently of RAD17, and can also be activated by other factors such as RPA-coated ssDNA [7], suggesting separable and complementary roles for ATR and RAD17.

In meiosis, Spo11-DSBs initiate homologous recombination and are therefore essential for the generation of crossovers (COs) between homologous chromosomes. In most organisms, including mammals and *S*. *cerevisiae*, the organism utilised in the work presented here, such recombination-dependent interactions facilitate homologue pairing during leptotene-zygotene, full alignment and connection via the synaptonemal complex at pachytene, and subsequent reductional chromosome segregation at the meiosis I nuclear division [8]. In *S*. *cerevisiae* meiosis, similar to the vegetative DDR pathway, the 9-1-1 clamp complex, its loader Rad24, and the Mre11-Rad50-Xrs2 complex act as damage sensors [9], with ssDNA, produced by the resection of Spo11-DSBs, activating Mec1 [10]. By sensing ongoing recombination activity and unrepaired DSBs, Mec1 acts as a molecular rheostat to modulate the progression of meiotic prophase via the Ndt80 transcription factor, and the Mek1 kinase, a paralogue of Rad53 (CHK2 in mammals; [10–13]). Due to this transient checkpoint activation, Mec1 is able to prolong the stage during which Spo11-DSB formation can occur [14], and both Mec1 and Rad24 promote CO formation and suppress ectopic (non-allelic) recombination [15,16]. The interaction between Mec1 and the 9-1-1 complex also contributes to other Mec1 functions such as phosphorylation of the meiosis-specific chromosome axis protein, Hop1 (HORMAD1/2 in mammals [17]), which is important for the maintenance of homologue bias and chiasma formation [18,19]. In mammals, ATR localises to meiotic chromosomes [20] and is a key regulator of meiotic events. ATR deletion in male mice causes fragmentation of the chromosome axis, and ATR is required for synapsis and loading of recombinases RAD51 and DMC1 at DSB sites [21,22]. In *Drosophila*, loss of Mei-41, the fly ATR orthologue, not only abrogates checkpoint signalling [23], but also alters the frequency and spatial distribution of COs [24]. In *Arabidopsis thaliana*, although *atr* mutants are fully fertile [25], ATR promotes meiotic recombination by regulating the deposition of DMC1 at DSBs [26].

Observations in *S. cerevisiae* indicate that Spo11 DSBs do not occur independently, but are instead subject to interference among the four chromatids (reviewed in [27]). This interference occurs in *cis* (adjacent, on the same chromatid [28]), and *trans* (between chromatids [29]). While both Mec1 and Tel1 are involved in DSB interference and the global suppression of Spo11 activity [28–30]— roles that appear conserved in other organisms [23,31]—a direct role for Rad24 is unclear [28]. CO events also display interference (reviewed in [32]), a process dependent upon the ‘ZMM’-family of proteins (reviewed in [33]). Rad24 is necessary for the efficient loading of ZMM proteins to meiotic chromosomes, and interacts physically with Zip3, independently of Mec1 [34], suggesting that Rad24, but not Mec1, may promote interfering CO formation. However, there has been relatively little investigation into the effects Mec1 and Rad24 may have upon CO interference.

In order to further establish the roles of Mec1 and Rad24 in the regulation of meiotic recombination, and to determine whether previously-observed roles can be extended genome-wide, we mapped meiotic recombination patterns at high resolution across the *S. cerevisiae* genome in *rad24Δ* and *mec1-mn (’meiotic null’)* backgrounds, revealing both similar and distinct roles for Mec1 and Rad24 in the regulation of meiotic DSB repair.

## Results

### Reduced spore viability in *mec1* and *rad24 mutants* can be alleviated by extending meiotic prophase or by deleting *MSH2*

To generate a genome-wide picture of meiotic recombination in wild type, *mec1* and *rad24* mutants, *Saccharomyces cerevisiae* SK1xS288c hybrid diploids (∼65,000 SNPs, ∼4,000 high confidence INDELs, ∼0.57% divergence) were sporulated, the resulting tetrad of spores separated and genomic DNA sequenced at an average depth of 44x. In strains with a deletion of the mismatch repair (MMR) protein, Msh2, additional heteroduplex (hDNA) tract information can be retrieved by allowing spores to undergo one round of mitotic division and separating them again to form an octad (post-meiotic segregation) prior to sequencing (Fig 1A; as described [35,36]). Sequences were aligned against SK1 and S288c reference genomes, and polymorphism information was subsequently used to identify regions of recombination using a custom pipeline (Fig S1 **and Methods**).

**Figure 1.**
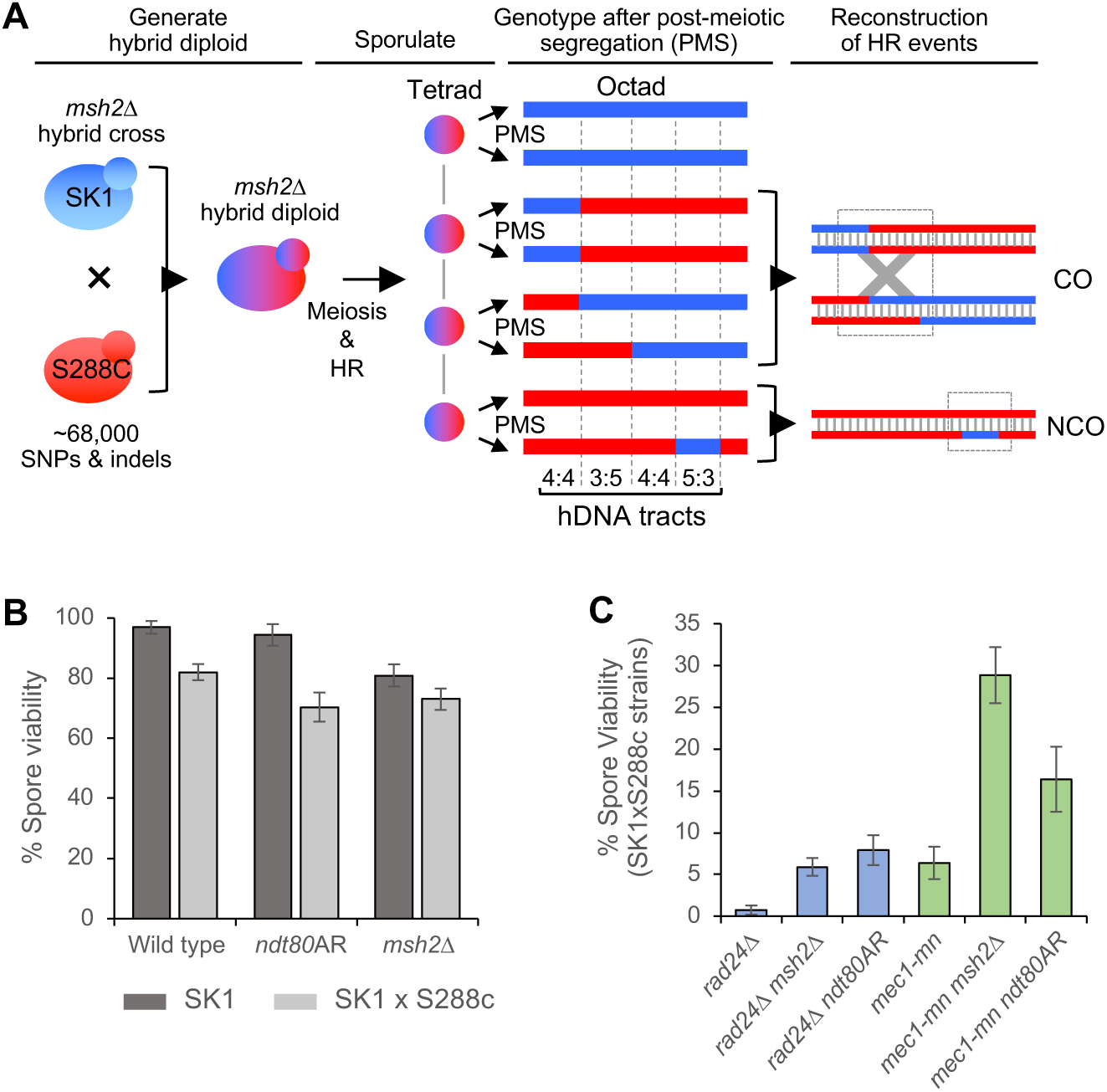
Overview of recombination analysis strategies. **A)** Strategy for whole-genome mapping of HR events in octads after post-meiotic segregation (PMS) of hDNA. For simplicity, a single chromosome is shown containing one CO and one NCO. Haploid cells from the S288c (red) and SK1 (blue) genetic backgrounds are crossed, producing a hybrid diploid containing many polymorphisms. The diploid undergoes meiotic recombination, forming CO and NCO events with associated hDNA tracts. These are left unrepaired in strains with a deletion of the mismatch repair protein Msh2, but are converted or restored in *MSH2* strains. At the conclusion of meiosis, the four chromatids are distributed among four ascospores. To preserve hDNA information, each spore is allowed to undergo one mitotic division, after which the mother and daughter cells are separated to produce eight haploid cell lines, the genomes of which are equivalent to the eight DNA strands involved in the meiotic recombination event. The eight genomes are sequenced to retrieve polymorphism information at each position, allowing precise hDNA reconstruction genome-wide. **B,C)** Spore viability is severely reduced in *rad24*Δ and *mec1-mn*, which is prohibitive to their analysis. To enable sequencing of all four meiotic products, spore viability is increased by prophase extension and *msh2*Δ deletion. Spore viability comparison for the indicated strains and non/hybrid backgrounds. Error bars indicate 95% confidence limits.

To avoid the influence of Mec1 inactivation during premeiotic growth, we used a conditional *MEC1* allele (*P*_*CLB2*_*-MEC1*, hereafter referred to as *mec1-mn* for ‘meiotic null’) that is only expressed during mitotic growth [14]. In *mec1-mn*, Mec1 protein becomes undetectable by three hours after meiotic induction [14], indicating that during the majority of DSB formation and repair there is expected to be little or no Mec1 protein present.

Accurate determination of recombination patterns by this method requires all spores to survive, hampering the analysis of genotypes with low spore viability (i.e. mutants such as *mec1* and *rad24* that perturb recombination and chromosome segregation). Spore viability is also reduced in hybrid strains compared to pure SK1 or S288c backgrounds (Fig 1B); potentially due to the effect sequence divergence has upon recombination [35,37,38]. Indeed, the already low spore viabilities of *rad24*Δ and *mec1-mn* mutants were exacerbated in the hybrid background (Fig 1C).

To overcome the barrier to analysis caused by low spore viability we explored three strategies. Firstly, we investigated the impact of deleting *SML1*, an inhibitor of ribonucleotide reductase that is normally inactivated by Mec1 in response to DNA damage, thereby increasing dNTP pool levels—promoting genetic fidelity and cell viability [39]. However, Sml1 was not previously known to have any role in meiosis, nor to affect the viability of a *rad24*Δ mutant. We found that *sml1*Δ increased the spore viability of *rad24*Δ hybrids ∼7 fold (to 4.7%; Fig S2A); nevertheless, only one four-spore viable tetrad was obtained from 196 tetrad dissections, limiting the practicality of this approach.

As a second strategy, we artificially lengthened meiotic prophase via an *NDT80* arrest-and-release (‘*ndt80AR*’) system that we previously described as a method to increase spore viability in *rad24*Δ mutants [14]. Normal prophase length is ∼4-5 hours, but the loss of the DNA damage response (DDR) checkpoint causes an early exit from meiotic prophase [12]. Spore viability was increased in both *rad24*Δ and *mec1-mn* mutants (Fig 1C) and was proportional to the length of time held in prophase (Fig S2B). Thus, the artificially extended prophase mimics, and somewhat extends, the checkpoint arrest that the DDR mutants lack [14]. Whilst extension of prophase significantly improved spore viability of *mec1* and *rad24* mutants, it did not greatly impact wild type spore viability (Fig 1B).

The final method explored to increase spore viability was deletion of *MSH2*, which was initially employed to preserve hDNA tract information for detailed recombination analysis [35,36]. Unexpectedly, deletion of *MSH2* increased spore viability in hybrid *rad24*Δ and *mec1-mn* strains (Fig 1C), despite causing a slight reduction in spore viability in wild type (Fig 1B). Whilst the precise mechanisms that increase spore viability are unclear, we consider it is likely due to the increased rates of recombination in hybrid backgrounds enabled by *msh2*Δ, and the removal of the role Msh2 plays in skewing recombination away from regions of high polymorphism density [35,37].

Based on the above findings, we reasoned that the cause of the low spore viability in *mec1-mn* and *rad24*Δ strains is primarily due to insufficient time for recombination to occur productively within the shortened prophase window of the DDR mutants—which can be rescued by either extending prophase, or by eliminating Msh2, a factor involved in rejecting mismatched strand invasion intermediates [40], something that will occur frequently in a highly polymorphic hybrid background.

### Recombination frequencies are altered in Mec1 and Rad24 mutants to different extents

By increasing spore viability to experimentally tractable levels, the *ndt80AR* and *msh2*Δ alleles provide an avenue to explore recombination patterns in DDR mutants for the first time. Nevertheless, before analysing the effect on recombination of losing Rad24 and Mec1 function, we first assessed the impact of the *ndt80AR* and *msh2*Δ alleles in an otherwise wild type background. In all *ndt80AR* strains, meiotic prophase was extended to 8 hours. Extending the length of meiotic prophase in this way increased CO and NCO frequencies ∼1.4-fold (Fig 2A), consistent with the persistent DSB signals observed in terminally-arrested *ndt80*Δ cells [41,42]. Inactivation of *MSH2* increased CO levels similarly (∼1.4-fold), but with a greater increase (∼2.9-fold) on NCOs, as has been observed previously [35]. The effects of *msh2*Δ are likely to be due to a combination of the increased visibility of NCOs in the absence of mismatch restoration, and the loss of mechanisms that inhibit recombination at sites of polymorphism in *MSH2*+ strains. No further increase in recombination was observed in the *ndt80AR msh2*Δ double strain (Fig 2A), suggesting that there may be an upper limit to recombination frequency in these backgrounds. Interestingly, extending prophase length, but not *MSH2* deletion, skews recombination towards subtelomeres (Fig S3A-B)—regions that are both less compacted as measured by Hi-C [43], and retain disproportionately high levels of DSB formation at pachytene [44].

**Figure 2.**
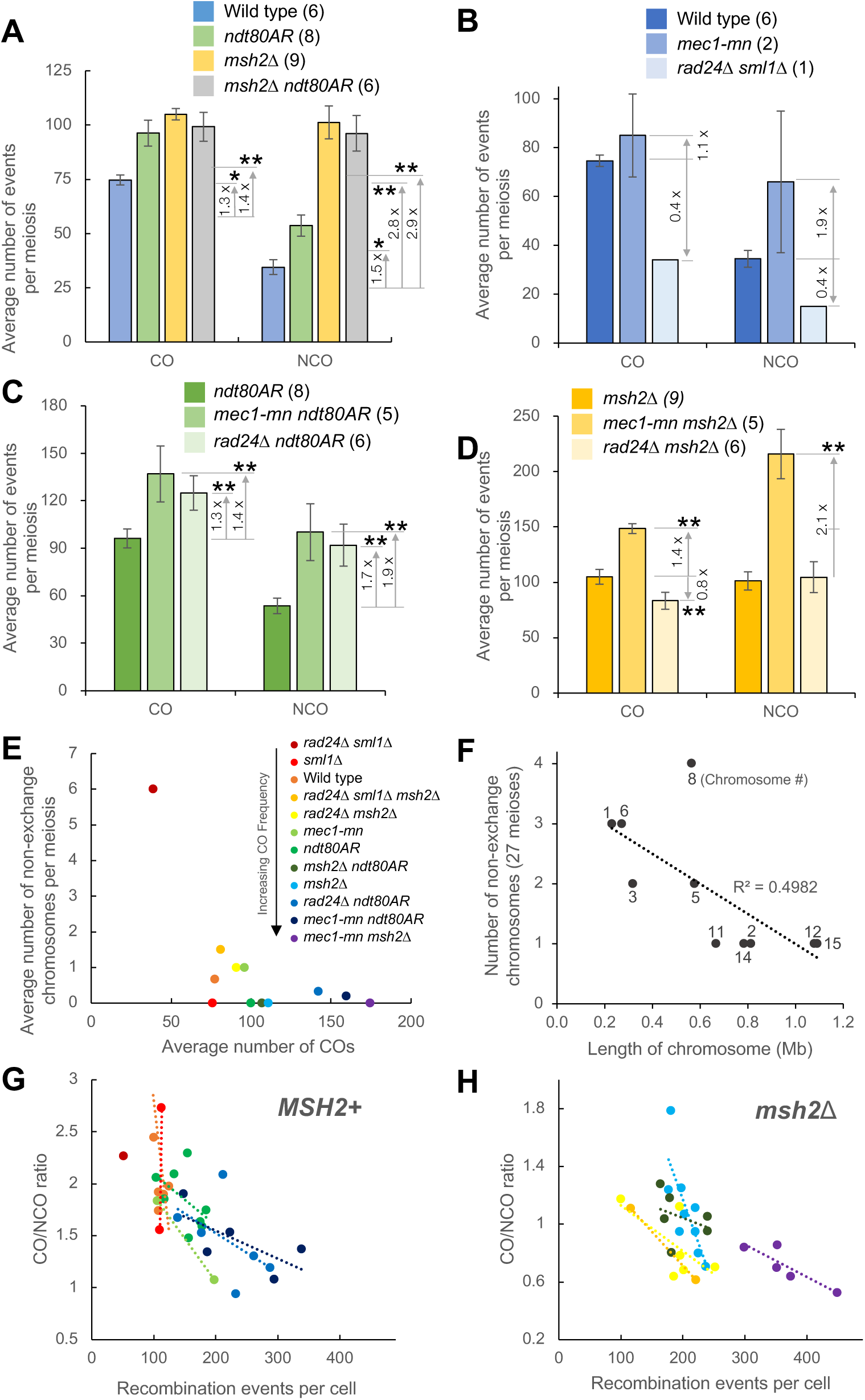
Meiotic recombination event counts and spatial regulation in the absence of Mec1 or Rad24. **A-D)** Recombination event frequencies in control strains **(A)** or in *rad24*Δ and *mec1-mn* with the genetic background **(B)** WT or *sml1*Δ; **(C)** *ndt80AR*; **(D)** *msh2*Δ. The mean number of CO and NCO events per octad/tetrad is shown (including both single and multi-DSB events). Error bars are standard error of the mean. Differences were tested by two-tailed T-test (* = P<0.05, ** = P<0.01). Significance values are not shown for comparisons between sets of *msh2*Δ/*MSH2* strains, due to the known increase in event visibility in *msh2*Δ strains. **E)** The average number of chromosomes without a crossover (CWCs) per octad/tetrad, plotted against the average number of crossovers. **F)** The relationship between the length of a chromosome and the number of times it was a non-exchange chromosome in 27 *mec1-mn* and *rad24*Δ mutant meiosis. **G,H)** The relationship between the recombination event count and the CO/NCO ratio in each cell. The plotted lines are linear models of all *MSH2* **(G)** or *msh2*Δ **(H)** strains. Colours indicate strains as in **(E)**.

Prior analysis indicated that *rad24*Δ mutants display a ∼10-20% reduction in global DSB formation compared to controls, thought to be because Mec1-mediated inhibition of Ndt80 is needed to allow sufficient time for wild type levels of DSBs to form [14]. By contrast, Mec1 has also been reported to suppress DSB formation via the process of *trans-*inhibition [29]. To investigate the relationship between DSB levels and global recombination rate, meiotic recombination was assayed in one *rad24*Δ *sml1*Δ and two *mec1-mn* tetrads (Fig 2B); five *mec1-mn ndt80AR* and six *rad24*Δ *ndt80AR* tetrads (Fig 2C); and five *mec1-mn msh2*Δ and six *rad24*Δ *msh2*Δ tetrads (Fig 2D).

The global recombination levels in *mec1-mn* were not significantly different to those of the wild type strain (Fig 2B). However, in both the *msh2*Δ and *ndt80AR* backgrounds, the *mec1-mn* mutant displays a ∼1.4x increase in CO formation and a ∼2.0x increase in NCO formation compared to the relevant controls (Fig 2C, D). The greater increase in NCOs than COs within *mec1-mn* strains suggests that CO homeostasis remains functional in the absence of *MEC1* activity. Interestingly, the enrichment in telomeric recombination, observed within *ndt80AR*, is suppressed within both *mec1-mn* and *rad24* mutants (Fig S3C-F), suggesting the abrogation of those processes that contribute to subtelomeric enrichment in DSB and recombination activity.

Increased recombination suggests that DSB formation may be increased above wild type levels in *mec1-mn* mutants—at least when the length of prophase is sufficient for this to happen— consistent with the role of Mec1 in DSB *trans*-inhibition [29]. Previously, we concluded that Mec1-dependent inhibition of Ndt80 also promotes meiotic DSB formation [14]. Our results here agree with this conclusion and suggest that delaying Ndt80 activation not only increases DSB frequency, but also the inter-homologue recombination rate.

In comparison to the *mec1-mn* mutant, we observed variable changes in recombination frequency in *rad24*Δ. Firstly, in the single viable *rad24*Δ *sml1*Δ tetrad, CO and NCO rates were reduced to less than half of the wild type values (Fig 2B). By contrast, the *rad24*Δ *ndt80AR* mutant displayed a significant increase in COs (∼1.3x) and NCOs (∼1.7x) compared to the *ndt80AR* control strain—a change not significantly different to that seen in *mec1-mn ndt80AR* (Fig 2C). The very similar recombination rate in both *ndt80AR* DDR mutant strains contrasts with the different rates observed between the strains with natural prophase length (Fig 2B). Although the inability to generate repeat datasets for *rad24*Δ *sml1*Δ hampers our statistical confidence, we tentatively conclude that the length of prophase has a greater positive impact on recombination rate in *rad24*Δ than in *mec1-mn*.

Loss of *RAD24* function in the *msh2*Δ strain led to no increase in recombination frequency (Fig 2D): COs were actually reduced ∼0.8-fold, whilst NCO frequencies were unchanged. These changes contrast strongly with both the impact of losing *MEC1* function in *msh2*Δ (above), and with the increased recombination observed in *rad24*Δ *ndt80AR* (Fig 2C). Such differing responses to loss of *MEC1* and *RAD24* in the *msh2*Δ background indicate that despite their known role in the DDR, Mec1 and Rad24 may function independently of one another under certain circumstances [34].

We reasoned that because *sml1Δ* does not compensate for the checkpoint or transcriptional functions of Mec1 [39], and because recombination rates were not significantly altered between wild type and *sml1*Δ, or between *rad24*Δ *sml1*Δ *msh2*Δ and *rad24*Δ *msh2*Δ (Fig S4), the level of recombination in *rad24*Δ *sml1*Δ may be similar to that of *rad24*Δ single mutants (which were precluded from our analysis because we were unable to obtain any four spore viable tetrads). This interpretation agrees with the prior observation that CO numbers are reduced in *rad24*Δ strains [15].

The reduction in recombination in *rad24*Δ *sml1*Δ is greater than anticipated considering the relatively mild reduction in DSB formation [14]. Notably, our recombination assay is only capable of detecting allelic recombination between homologues, suggesting that the *rad24*Δ mutant may have an increase in inter-sister recombination [15,34], which would be undetectable. Surprisingly however, despite Mec1’s known role in ensuring homologue bias ([18], observable rates of recombination were not reduced in *mec1-mn*. We continue to explore differences and similarities caused by loss of Mec1 and Rad24 activity in meiotic recombination throughout this study.

### Increased occurrence of non-exchange chromosomes in DDR mutants

To aid the correct disjunction of chromosomes during meiosis I, it is important that each chromosome receives at least one CO per homolog pair—a process known as CO assurance [45,46]. To investigate whether this aspect of regulation is intact within Mec1 and Rad24 mutants, the frequency of non-exchange chromosomes was assessed (Fig 2E, Table S1). Strikingly, the single four spore viable *rad24*Δ *sml1*Δ tetrad contained six chromosomes with no detectable CO or NCO (Fig 2E). Additionally, half of the twelve *rad24*Δ *ndt80AR* or *rad24*Δ *msh2*Δ tetrads lacked a CO on at least one chromosome (Fig 2E). In contrast, only one out of ten *mec1-mn ndt80AR* or *mec1-mn msh2*Δ tetrads contained a chromosome without a CO, suggesting that the mechanisms contributing to CO assurance are less dependent on Mec1 than on Rad24, which may help explain why *mec1-mn* mutants have a higher spore viability than *rad24*Δ.

The occurrence of non-exchange chromosomes tends to anti-correlate with CO counts (Fig 2E), and with chromosome size (Fig 2F), suggesting that a sufficiently high CO frequency will help to ensure that each chromosome receives a CO. However, some of the *rad24*Δ *ndt80AR* and *mec1-mn ndt80AR* tetrads—which have higher CO counts than their controls—nonetheless displayed some nonexchange chromosomes, indicating that CO frequency is not the only contributor to gross CO distribution.

To examine CO homeostasis—the process that ensures a stable CO frequency despite variability in DSB number [47]—we plotted the ratio of CO:NCO formation against the total number of detectable recombination events for *MSH2*^+^ and *msh2*Δ strains (Fig 2G, 2H respectively). CO:NCO ratios cannot be compared between *MSH2*^+^ and *msh2*Δ datasets due to the increased visibility of NCOs in a *msh2*Δ background. CO homeostasis, shown as a negative correlation on these plots, is present within all mapped strains except wild type and those in an *sml1*Δ background—suggesting that neither Rad24 or Mec1 has an appreciable role in this process. The absence of detectable homeostasis within wild type is likely due to an unusually low NCO detection rate (see Fig 1A).

### Loss of *RAD24* and *MEC1* function increases the frequency of recombination events initiated by more than one DSB

Spo11-DSBs and the resulting CO events are spread across the genome more evenly than expected by chance due to the combined processes of DSB interference [28,29] and CO interference [27,32]. Previously, Mec1 has been implicated in the process of *trans* DSB interference [29], whereas Rad24 is likely to have an indirect influence on CO distribution given its importance in helping to load the pro-class I CO factor, Zip3 [34]. To investigate the role of Mec1 and Rad24 in DSB interference, we assessed the frequency of recombination events that were compatible with initiation by more than one DSB, something that is expected to be infrequent in wild type cells but frequent in cells lacking interference [28]. Such ‘multi-DSB’ events, are defined here as recombination events that can be explained by the formation of two or more separate Spo11-DSBs arising within 1.5 kb, and which are initiated on independent chromatids (Fig 3A, Fig S5).

**Figure 3.**
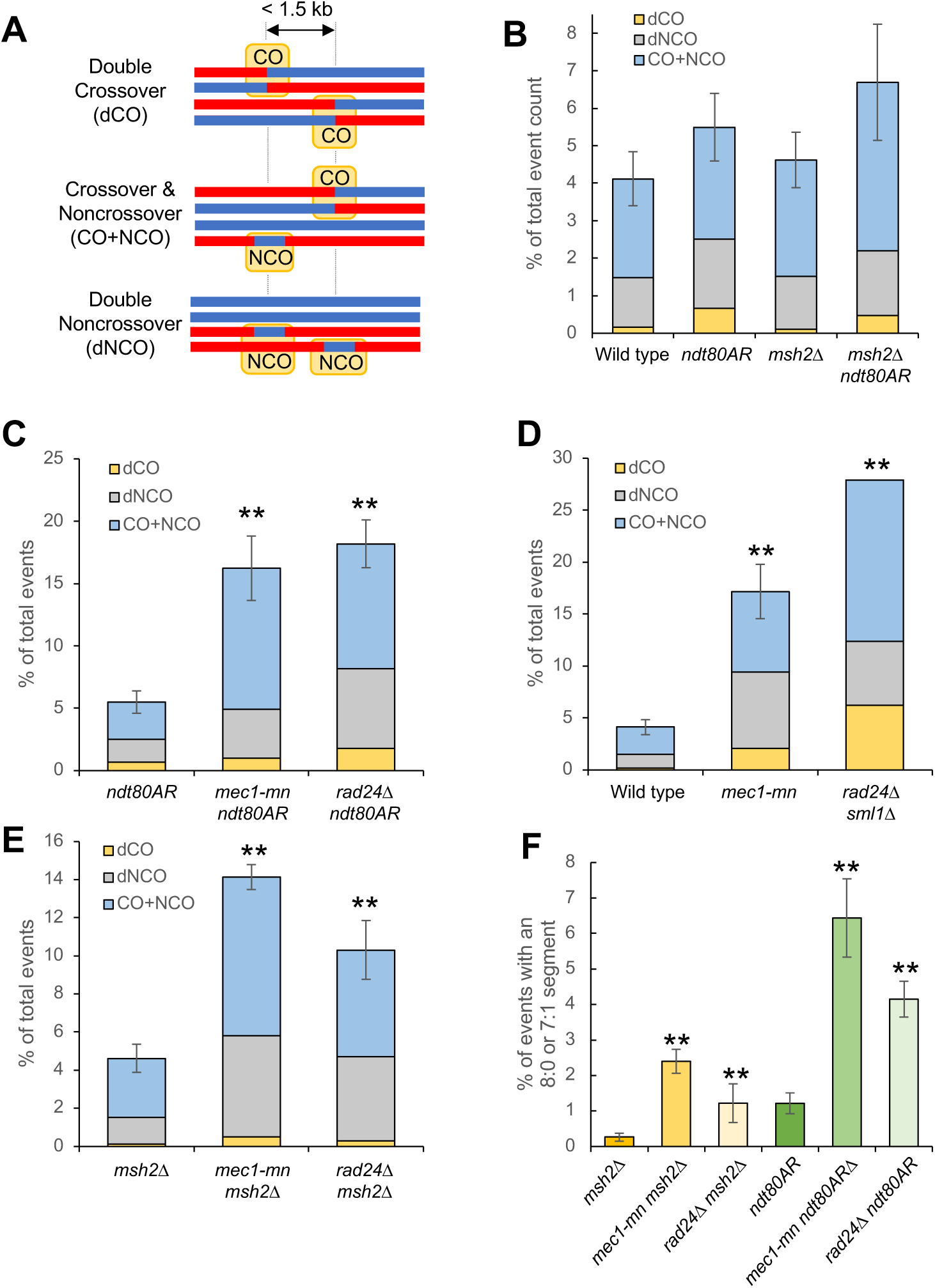
Meiotic recombination frequency and spatial regulation in the absence of Mec1 or Rad24. **A)** Simple depiction of the multiple-DSB event categories. **B-E)** Proportion of recombination events that are thought to result from multiple DSBs in **(B)** control strains, **(C-E)** *rad24*Δ and *mec1-mn* with the genetic background **(C)** WT or *sml1*Δ; **(D)** *ndt80AR*; **(E)** *msh2*Δ. The average proportion of multiple-DSB events per octad/tetrad is shown. Error bars are standard error of the mean. Differences in the proportion of multi-DSB events in each mutant and in the appropriate reference strain were tested by Fishers exact test (* = P<0.05, ** = P<0.01). **F)** The average number of directly overlapping events per octad/tetrad, defined as containing a segment of 8:0 or 7:1 segregation. All such events are also considered to be multi-DSB events of some kind. Differences were tested by Fishers exact test (* = P<0.05, ** = P<0.01). Differences were not tested between *MSH2* and *msh2*Δ backgrounds because 7:1 segments are not generated in *MSH2* backgrounds.

Potentially, a second DNA break could be formed by something other than Spo11 during the repair of a first DSB. For example, the endonuclease Mlh1-Mlh3, which has CO resolution activity in meiosis, also exhibits DNA cleavage activity (Manhart *et al*., 2017). To reduce the potential impact of such factors upon the analysis, the classification as a multi-DSB event is highly conservative in nature. Specifically, double COs (dCOs) are only considered when they affect all four chromatids in a double reciprocal exchange. Although it is possible for a dCO to involve two or three chromatids, the resulting patterns cannot be unambiguously distinguished and so are discarded from our analysis. Additionally, a CO+NCO cluster is only included in the analysis when the NCO falls on a third chromatid. Double NCOs (dNCOs) are only included when occurring on two sister chromatids or on two homologues with perfect overlap (Fig 3A and Fig S5 A-D). Due to the stringency of this analysis, it is likely that the true number of multi-DSB events is greater than presented.

Multi-DSB events were relatively infrequent in wild type cells (∼5% of events), and were not significantly increased by *msh2*Δ or *ndt80AR*-mediated prophase extension (Fig 3B). By contrast, both *mec1-mn* and *rad24*Δ strains, in all genetic backgrounds assayed, have a significant (∼3-fold) increase in the formation of multi-DSB events compared to the relevant control (Fig 3 C-E). Thus, the spatial regulation of DSB formation seems to be regulated to a similar degree by both Mec1 and Rad24. Within multi-DSB events, the proportions of dCO, dNCO and CO+NCO events were not significantly altered between strains (Fig 3 B-E).

Events seemingly formed by multiple DSBs on different chromatids could potentially arise from multi-strand invasions, whereby the invading filament sequentially repairs using multiple chromatids [48]. To eliminate the possible influence of multi-strand invasions on our multi-DSB event calculations, we looked specifically at the formation of multi-DSB events containing a segregation pattern of 8:0 or 7:1 (Fig S5 E-G). Because conversion on two sister chromatids at the same genetic locus is necessary to form these patterns, they are strong indicators of multi-DSBs, and cannot be easily explained by multi-strand invasion.

The occurrence of events with 8:0 and 7:1 segregation patterns (“*trans*-DSB” events) is significantly increased in both *mec1-mn* and *rad24*Δ mutants compared to controls (Fig 3F; Table S1). These observations support and extend the known role of Mec1 in promoting *trans*-DSB interference [29], and suggest a similar role for Rad24. Taken together, these results highlight a failure of spatial regulation to control DSB formation within Mec1 and Rad24 mutants, marked by a loss of DSB interference in *trans* and an increased occurrence of nonexchange chromosomes, which impacts chromosome segregation and spore survival.

### Recombination tract lengths are increased in DDR checkpoint mutants

The number of genetic markers involved in a recombination event, and the total genomic distance that they span (‘event length’), may be influenced by dHJ branch migration, joint molecule migration, additional DNA strand breakage during repair [36], or by the distance of DSB resection, which averages ∼1.5 kb in wild type cells [49,50]. It is currently unclear to what extent each of these factors contribute to the final event length, or how DDR factors may contribute. Notably, hyper-resection at Spo11-DSBs has been observed in DDR mutants such as *rad24*Δ, *rad17*Δ and *mec1-mn* [10,14,16]; K. Wardell & M. Neale, unpub. obs.), suggesting that if DSB resection is a significant contributor to the overall amount of DNA involved in each recombination event, tract lengths should be increased in *rad24*Δ and *mec1-mn* strains.

Because event lengths are affected by local marker density, estimates were made using the midpoint between the markers flanking each event (Fig 4A). To avoid the complexities inherent within overlapping multi-DSB events, only unambiguous single-DSB events were considered. To compare between samples, median recombination event lengths, along with upper and lower quartile values, were calculated for each genotype (Fig 4B-G, Fig S6).

**Figure 4.**
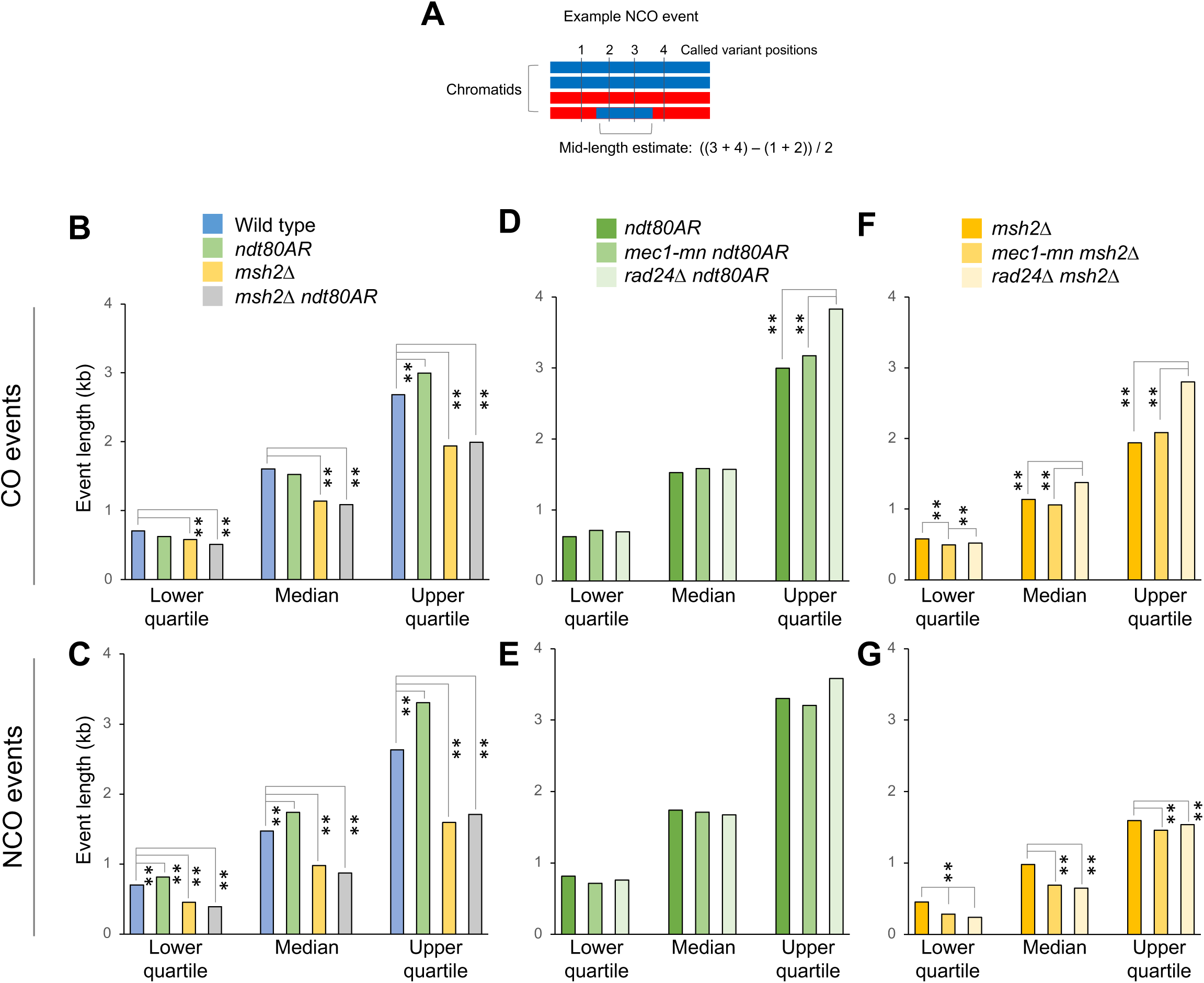
Meiotic recombination event tract lengths in the absence of Mec1 or Rad24. **A)** Simple depiction of event mid-length calculation. The mid-length estimate is defined as the distance between the midpoints of the intermarker intervals surrounding the recombination event. The lengths of multi-DSB events are omitted. **B-G)** The median, upper and lower quartile values for the mid-lengths of the strand transfers associated with CO **(B,D,F)** and NCO **(C,E,G)** events in the indicated strains. Statistical differences between strains are indicated (Wilcoxon test).

Within control strains, *MSH2* deletion led to a significant decrease in event length, which was most pronounced for NCOs (Fig 4B,C), revealing that mismatch correction leads to gene conversion of markers beyond just the regions of hDNA generated directly by the recombination process. By contrast, extending prophase length via *ndt80AR* moderately increased event lengths, an effect that was most pronounced in the *MSH2*+ background (Fig 4B,C)—perhaps due to persistent MMR activity, and/or increased opportunity for branch migration of persistent recombination intermediates present in prophase-arrested cells [42].

Loss of Rad24 or Mec1 activity had complex effects on recombination event lengths (Fig 4D-G). Whilst median lengths were broadly unaffected, longer recombination events were disproportionately extended—with the upper quartile event length values increased up to ∼1.4-fold in DDR mutants compared to controls (Fig 4D-G). These increases in event length may indicate that the hyper-resection evident in DDR mutants via physical analysis has an influence in the final pattern of recombination. It is also possible that DDR components influence other aspects of recombination, such as JM migration and/or stability, leading to longer tracts of genetic change in the haploid progeny. Intriguingly, NCO event lengths in the *msh2*Δ background are significantly shorter in both DDR mutants (Fig 4G). Thus it is possible that DDR components increase the stability of recombination intermediates—perhaps related to their known role in suppressing ectopic recombination [15,16,S. Gray & M. Neale unpub. obs.]— thereby leading to shorter tracts of repair synthesis when mutated. Importantly, polymorphism density around NCO events was unchanged in the absence of Rad24 and Mec1 (∼7 variants per kb), indicating that the reported changes in event length are not technical consequences of a redistribution of NCO events towards more highly polymorphic areas, which may have led to a more accurate estimate of event length.

### Detection of heteroduplex DNA in very low spore viability strains

Genome-wide analysis of meiotic recombination in *msh2*Δ octads permits the detection of not only hDNA, but also the determination of strand polarity, which enables a more detailed inference of recombination mechanism [35,36]. Therefore, in order to investigate the role of DDR components in specific pathways of recombination, HR strand transfer patterns were explored in more detail within *mec1-mn msh2*Δ and *rad24*Δ *msh2*Δ meioses compared to controls, by computationally and manually sorting into categories as previously described [36]. Example events from each category are shown in (Fig S7 and S8).

Due to low spore viability, it was impractical to directly generate and analyse octads in *rad24*Δ *msh2*Δ and *mec1-mn msh2*Δ strains. Instead, heteroduplex information was recovered from *msh2*Δ tetrads, by converting markers displaying ∼50% coverage of each genotype (“heteroduplex” calls) into one copy of each parental SNP (See Methods) to generate pseudo-octads (Fig 5A). Although this method allows the isolation of hDNA patches, information on strand polarity is not retrieved. To additionally recover strand polarity information, post-meiotic haploid cells were streaked onto rich media, and a single colony sequenced at high depth (Fig 5A). Due to postmeiotic segregation, such clones will retain the complete haplotype information of one of the strands present in the mixed reads from the *msh2*Δ tetrad. By comparing this haplotype to the mixed heteroduplex reads, the haplotype of the other member of the pair can also be constructed (Michael Lichten, pers. comm.). Haplotype re-sequencing by this method was performed for a limited number of *msh2*Δ *rad24*Δ and *msh2*Δ *mec1-mn* meioses (Fig 5B,C).

**Figure 5.**
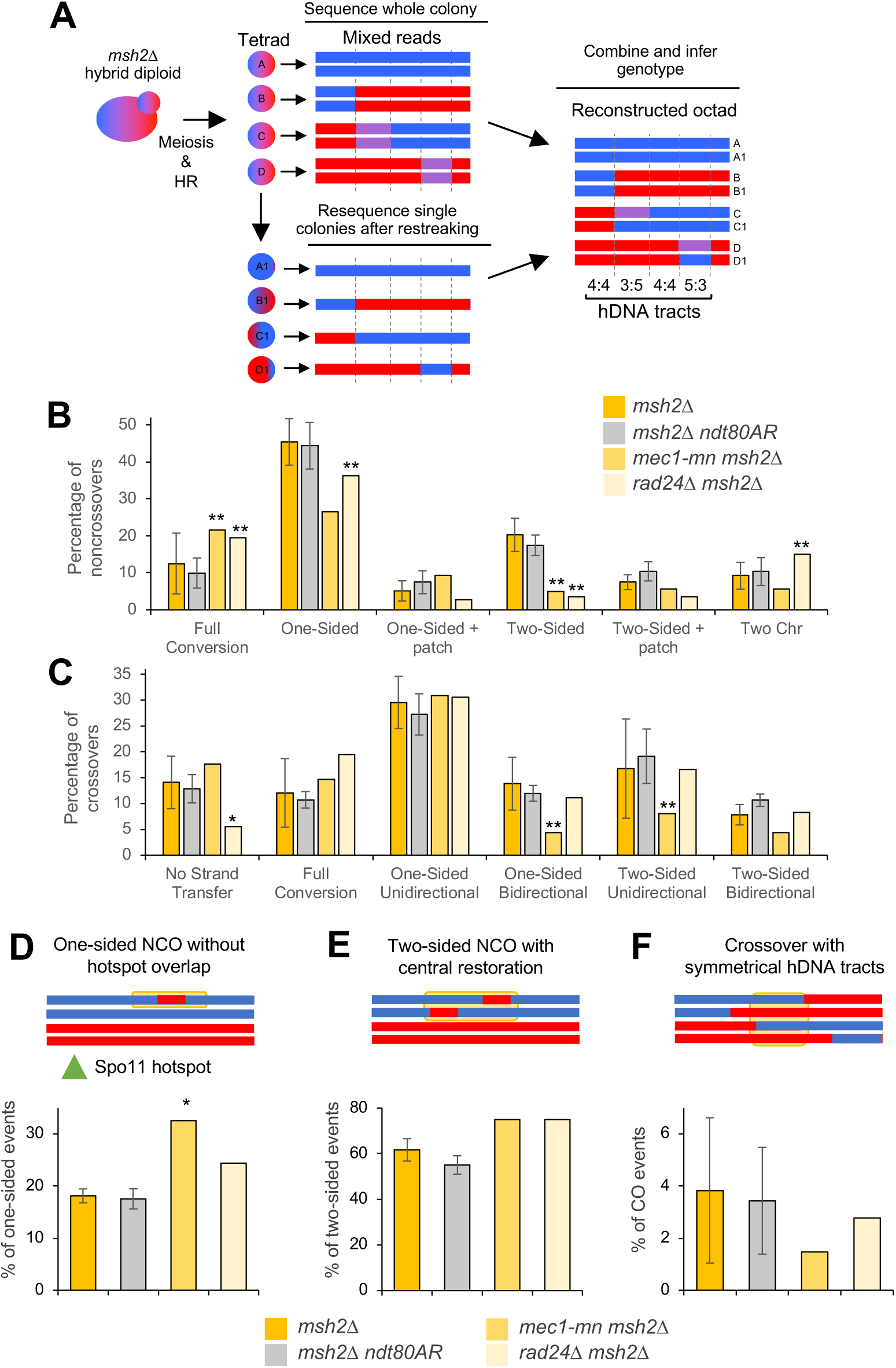
Subcategorization of single-DSB events based on hDNA patterns. **A)** Strategy for the retrieval of hDNA in low-viability backgrounds. *msh2*Δ strains are mated and sporulated as in Figure 1A, and an entire sectored colony is sequenced, containing mixed reads at positions of hDNA. Additionally, a restreaked single colony is sequenced; the genotype of this colony at each position containing mixed reads allows the genotype of the other strand to be inferred. **B,C)** Crossover **(B)** and noncrossover **(C)** categories in *msh2*Δ, *msh2*Δ *ndt80AR, rad24*Δ *msh2*Δ and *mec1-mn msh2*Δ. All error bars are standard error of the mean. Stars indicate when the proportion of multi-DSB events is significantly different in the mutant and in the *msh2*Δ reference strain (one star, p < 0.05; two stars, p < 0.01, Fishers exact test). Only the *rad24*Δ *msh2*Δ and *mec1-mn msh2*Δ tetrads that were able to undergo detection of hDNA polarity were used (*mec1-mn msh2*Δ #3 and *rad24*Δ *msh2*Δ #6 in Table S1). Crossover and noncrossover events were classified according to their strand transfer patterns, illustrated in Fig S5 and S6. The percentage of each category over the total number of crossovers or noncrossovers is shown. Complex event types (Figure S4) contribute to the total number of COs and NCOs and thus affect percentages, but are not plotted again here. All categories are mutually exclusive. **D,E,F)** Proportion of events that appear to have involved template switching during repair. A simple example of each category is given, with the yellow box highlighting the region thought to have undergone sister chromatid invasion. **D)** One-sided NCOs that have occurred without overlapping a hotspot. **E)** Two-sided NCOs with a central patch of 4:4 restoration. **F)** Crossovers with symmetrical hDNA tracts.

Recombination events in nine control *msh2*Δ and six *msh2Δ ndt80AR* octads, were categorised and compared with the reconstructed *mec1-mn msh2*Δ and *rad24*Δ *msh2*Δ octads. Notably, the proportions of CO (Fig 5B) and NCO (Fig 5C) events detected in each category were not significantly altered between *msh2*Δ and *msh2*Δ *ndt80AR* octads, indicating that the prophase arrest does not impact the way in which DSBs are repaired. In the DDR mutants, the frequency of one-sided and two-sided NCOs was significantly reduced, whereas full conversion NCOs were elevated (Fig 5B). One- and two-sided NCOs are thought to be produced by the SDSA pathway [36]. Full conversion patches (regions of 6:2 segregation) may be produced by single-stranded nicking during repair, or via the repair of gaps generated by adjacent Spo11-DSBs, suggesting that one or both processes are subject to regulation by Mec1 and Rad24, the latter of which is consistent with loss of *trans*-DSB interference.

Interestingly, *rad24*Δ *msh2*Δ also displayed a small but significant increase in NCOs with changes on two homologous chromatids (Fig 5B; ‘Two Chr’)—indicative of an increase in NCO formation by dHJ resolution—something that was not observed in *mec1-mn msh2*Δ. Because these types of event are also more frequent in a *zip3*Δ [48], our results are consistent with Rad24—independently of Mec1—having a role in the loading of ZMM proteins at the sites of future CO events [34]. Specifically, the loss of ZMM loading in *rad24*Δ may mean that an increased number of joint molecules that would otherwise become class I COs, are instead resolved by class II factors such as Mus81, which is thought to be unbiased in its production of CO and NCO events. Consistent with this view, the CO:NCO ratio was lower in *rad24*Δ *msh2*Δ than the *msh2*Δ control (see Fig 2H).

Loss of Mec1 and Rad24 function also had slightly different effects on hDNA patterns associated with COs. The *mec1-mn msh2*Δ octad displayed a significant reduction in the frequency of one-sided bidirectional COs and two-sided unidirectional COs than both the *msh2*Δ control and the *rad24*Δ *msh2*Δ octad (Fig 5C). Bidirectional CO patterns, and some two-sided unidirectional COs with *trans* hDNA on one chromatid, are thought to be produced by dHJ migration [36]—suggesting that dHJ migration is reduced in *mec1-mn msh2*Δ. The only CO category altered in frequency in the *rad24*Δ *msh2*Δ octad was a decrease of COs with no associated strand transfer pattern (Fig 5C). COs with no strand transfer patterns are those occurring in regions of low marker density, and so have no associated hDNA information. Potentially, this could suggest a preference for COs in regions of higher marker density in *rad24*Δ.

### The role of Mec1 and Rad24 in maintaining Inter-homolog bias

Mec1, along with Tel1, promotes inter-homolog recombination in meiosis via the phosphorylation of Hop1 [18]. We were thus interested to investigate whether recombination patterns in *rad24*Δ and *mec1-mn* mutants displayed evidence of reduced homologue bias. While it is not possible to directly observe inter-sister recombination using our assay, we reasoned that alterations in the frequency of template switching (initial repair with the sister chromatid, then switching to repair with the homolog) in DDR mutants may suggest changes in the strength of inter-homologue bias.

Events containing patches of putative template switching were categorised by the presence of unconverted markers in recombination segments expected to be converted during normal DSB repair (Fig 5D-F) [36]. Surprisingly, only one class was altered: the proportion of one-sided NCOs that occur away from a Spo11-hotspot, and this was only increased in *mec1-mn* (P<0.05) (Fig 5D), suggesting either that there is redundancy in Hop1 activation and the promotion of inter-homologue bias, or that our analysis is not sensitive enough to detect major changes. Notably, *mec1-mn* strains display the greatest frequency of recombination events (up to ∼360 in total in *mec1-mn msh2*Δ) of any strain analysed—indicating that, at least in those meiosis analysed here, interhomologue recombination is not impeded.

### The global strength of crossover interference is reduced in *MEC1* and *RAD24* mutants

An important aspect of spatial regulation in meiosis is CO interference, a process whereby COs are spaced more evenly than would be expected from a random distribution [32]. While the DDR component, Tel1, has been implicated in CO interference [51], roles for Mec1 and Rad24 are unclear, and may be distinct [34].

To examine the role of Mec1 and Rad24 in CO interference, inter-CO distances (ICDs)—the distances (in bp) between successive COs along each chromosome—were calculated for *mec1-mn* and *rad24*Δ strains in both *msh2*Δ and *ndt80AR* backgrounds, arranged in order from smallest to largest and plotted as cumulative fraction of total CO events against ICD size (a cumulative distribution function, CDF, plot; Fig 6A-D). ICDs are further transformed (**Methods**) to enable direct comparisons between samples of different sizes [Cooper et al. 2018]. For completeness, untransformed plots are presented in (Fig S9).

**Figure 6.**
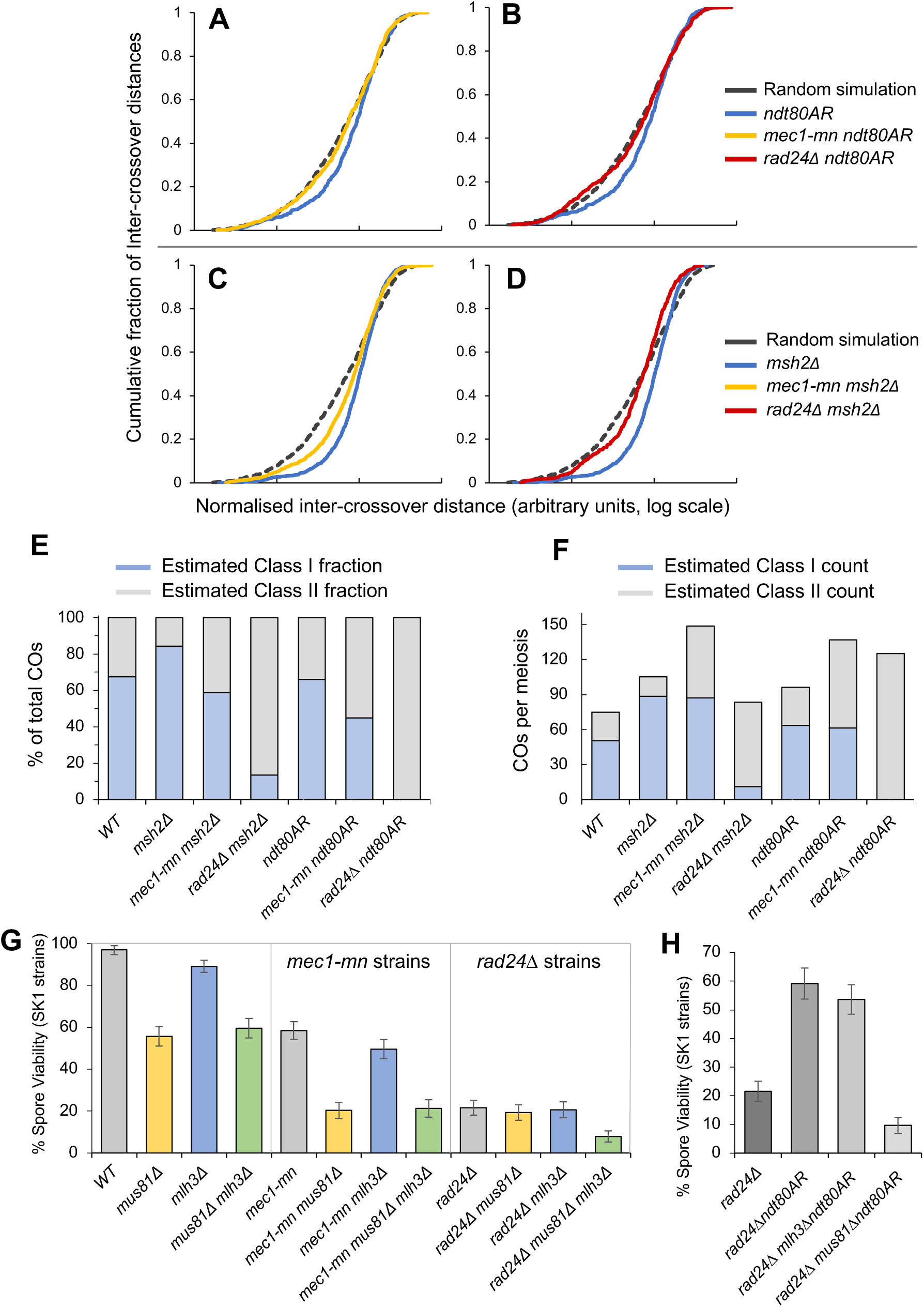
Inter-CO distance in Mec1 and Rad24 mutants compared to control strains. **A-D)** Inter-CO distances (ICDs) were calculated for the indicated strains, rank ordered, and plotted as cumulative fraction of the total CO count against inter-CO distance (ICD) on a log scale. For comparison, simulated datasets were generated where the same number of CO events detected in each strain were distributed randomly across the genome. All ICD values were transformed to enable inter-strain comparison of distributions (see **Methods** for details; [37]). **E, F)** Predicted Class I/Class II CO proportions **(E)** and frequencies **(F)**, based on a computational model (**Methods**). For each genotype, a mixture model was used to extract two component distributions, random and interfering, from the distribution of inter-CO event distances. The estimated proportion of CO events in each component can be applied to the actual event counts to estimate how many Class I (interfering) and Class II (random) COs there are in each genotype. The only strain in which two separate distributions could not be extracted was *rad24*Δ *ndt80AR*, which behaves as if there is a single, completely random component. **G, H)** Impact of the loss of CO resolution factors Mlh3 (Class I COs) and Mus81 (Class II COs) on the spore viability of *rad24*Δ and *mec1-mn* SK1 strains. Spore viability comparison between **G)** WT, *rad24Δ* and *mec1-md* SK1 strains with or without Mlh3 or Mus81; **H)** *rad24Δ* SK1 strains with or without 8 hours of *ndt80AR* and Mus81/Mlh3. Error bars are 95% confidence limits.

In order to visualise CO interference, experimental ICD distributions were compared to ICDs obtained from a simulation in which all COs are randomly placed. In all control strains, experimental ICDs display a distinctive sigmoidal curve that is shifted to the right relative to the random simulation, indicative of more even spacing between CO events (i.e. CO interference) (Fig S9A-D). Notably, loss of Msh2 activity leads to a more pronounced skew in this curve (Fig S9D), indicative of stronger interference, a phenomenon described in detail elsewhere [37]. By contrast, prophase extension, mediated via the *ndt80AR* allele, has no effect on ICD distribution (Fig S9C).

Loss of either Mec1 or Rad24 activity led to a shift in ICD distribution towards that of the random simulation (Fig 6A-D), consistent with a reduction in the global strength of CO interference. This shift is most pronounced in the *rad24*Δ *ndt80AR* strain, where the ICD plot became statistically indistinguishable from that of the random distribution (Fig 6B). Nevertheless, because the *rad24*Δ *msh2*Δ ICD distribution retained a small deviation from the random simulation (Fig 6D), we infer that CO interference is not entirely abolished in *rad24*Δ mutants—something that is only revealed by the overall increase in the global strength of CO interference when Msh2 activity is abolished.

Comparing normalised ICD plots suggests that CO interference is impacted slightly more severely by loss of Rad24 than Mec1 (Fig 6A-D). Consistent with this difference, Rad24 has been shown to promote Mec1-independent loading of ZMM proteins at sites of future interfering COs (Shinohara *et al*., 2015), suggesting that the different impact of *RAD24* and *MEC1* mutation on both CO distribution (Fig 6C-F) and spore viability (Fig 1C) may arise via alterations in the fraction of class I (interfering) COs.

### Reduced CO interference can be explained by a change in the ratio of Class I:Class II Cos

The pattern of meiotic COs observed within genome-wide maps is a composite of two overlapping distributions: a non-random pattern created by the interfering class I COs mixed with a randomly distributed pattern of non-interfering class II COs. Random (class II) and non-random (class I) components of mixed ICD distributions can be isolated using a mixture model algorithm [37]—a computational method that we have used to determine that the global increase in CO interference present in strains lacking Msh2 is explained by an increase in class I CO formation, rather than an alteration in the strength of interference between class I COs [37].

To investigate the role of Mec1 and Rad24 in class I CO formation, best-fit two-component distributions were obtained, generating estimated proportions and counts of class I and class II COs in each strain (Fig 6E-F; Table S2). A two-component solution was not generated for *rad24*Δ *ndt80AR*, because this strain is instead best fit by a single, random component. Kolmogorov-Smirnov tests provide statistical confidence that two-component distributions are good descriptions of the observed data (Table S2), something that was also evident by visual comparison between observed and estimated ICD plots (Fig S10).

Loss of Mec1 activity led to a ∼0.7-fold decrease in the estimated fraction of class I COs relative to control *msh2*Δ or *ndt80AR* strains (Fig 6E), whereas the impact of losing Rad24 activity was much more pronounced (∼0.16-fold in *rad24*Δ *msh2*Δ; a class I CO component was not detected within the ICD distribution of *rad24*Δ *ndt80AR*; Fig 6E). Mixture modelling also enabled estimated class I proportions to be converted to estimated class I CO frequencies per meiosis (Fig 6F). Remarkably, loss of Mec1 activity leads to no alteration in the overall estimated frequency of class I COs per meiosis (∼88 per meiosis in the *msh2*Δ background, and ∼62 per meiosis when prophase was extended via the *ndt80AR* allele; Fig 6F). Indeed, the large increase in COs evident in *mec1-mn* mutants appears to arise solely from a ∼4-fold increase in class II COs (Fig 6F). By contrast, *rad24*Δ *msh2*Δ retained an estimated average of only ∼11 class I COs per meiosis (Fig 6F), less than one per bivalent, a defect that we assume is even more pronounced in the presence of *MSH2* where class I CO frequencies are generally lower (Fig 6F).

These results suggest that the apparent similarity in the reduction of CO interference strength in *mec1* and *rad24* mutants actually arises for two completely separate reasons: (i) increased formation of non-interfering class II COs in *mec1-mn*, likely owing to excessive DSB formation upon loss of trans-DSB interference and (ii) by failing to efficiently load ZMM components, *RAD24* mutants have a disproportionate decrease in COs that are subject to interference (see also, **Discussion**). Importantly, mixture modelling reveals that the estimated strength of class I CO interference (as described by the gamma distribution parameter, α) were broadly similar in all strains (Table S2), suggesting that the overall mechanism of CO interference itself is largely independent of DDR activity.

### Spore viability is reduced disproportionately by the absence of Mus81 in checkpoint mutants compared to wild type strains

ZMM-dependent, class I, COs are resolved by the Mlh1-Mlh3 complex [45,52,53] whereas class II CO formation requires the nuclease activity of Mus81/Mms4, Yen1 or Slx1-Slx4 [53–58]. To further investigate the recombination pathways that are functioning when *RAD24* and *MEC1* are absent, we compared spore viability in a variety of class I and class II CO mutants in combination with *rad24*Δ or *mec1-mn* (Fig 6G). Loss of Mus81 activity led to a reduction in spore viability that was twice as severe in *mec1-mn* strains (from ∼58% to ∼20%; ∼0.3-fold) as it was in wild type (from ∼97% to 56%; ∼0.6-fold; Fig 6G), in agreement with the elevated frequency of class II COs arising in *mec1-mn* strains (Fig 6F). By contrast, loss of Mlh3 activity caused a similar reduction in spore viability in *mec1-mn* strains (from ∼58% to ∼50%; ∼0.9-fold) as in wild type (from ∼97% to 89%; ∼0.9-fold; Fig 6G), consistent with class I CO frequency being very similar in the presence and absence of Mec1 (Fig 6F). Loss of both Mus81 and Mlh3 activity gave no additional reduction in spore viability in either *mec1-mn* strain relative to the *mec1-mn mus81*Δ double mutant (Fig 6G), suggesting that the remaining DSB repair pathways are not influenced by the activity of Mec1.

To our surprise, and contrary to the above observations made in wild type or *mec1-mn* strains, individual loss of either the Mus81 or Mlh3 pathways had little impact on spore viability in the *rad24*Δ mutant (Fig 6G), which was unexpected given the mixture modelling estimations that suggest a very high proportion of class II COs in *rad24*Δ (Fig 6E, F). Nevertheless, combined loss of both CO pathways, resulted in a similar proportional decrease in spore viability in the absence of Rad24 (from ∼20% to ∼8%; ∼0.4-fold; Fig 6G) as reported above for wild type and *mec1-mn*.

Importantly, the *rad24*Δ strains in which recombination patterns were analysed utilised *msh2*Δ and *ndt80AR* alleles to increase spore viability—which arises, at least in part, by increasing the frequency of COs per meiosis (Fig 2B-D), but which may then place an extra burden on the recombination machinery. To test this idea we examined the impact on spore viability of removing either Mlh3 or Mus81 activity in the prophase-extended *rad24*Δ *ndt80AR* strain (Fig 6H). As predicted from the mixture modelling results, the spore viability increases in *rad24*Δ strains mediated by the *ndt80AR* prophase extension, were largely independent of Mlh3 activity (∼59% to ∼54%; ∼0.9-fold change), but very strongly dependent on the activity of Mus81 (∼59% to ∼10%; ∼0.2-fold change).

## Discussion

### Separable roles of Mec1 and Rad24 in regulating meiotic recombination

We report here the impact that the DNA damage response proteins Mec1 and Rad24 have on many aspects of meiotic recombination (Fig 7). Rad24 is known to be important for activating Mec1 via its role in loading the 9-1-1 clamp in response to ssDNA produced by the resection of DSBs [2]. Thus, as examined here, it is not surprising that in some aspects of meiotic recombination the effects of *rad24*Δ are similar to those of *mec1-mn*. For example, both Mec1 and Rad24 limit total recombination frequency (Fig 2B-D), and prevent the formation of multi-DSB events (Fig 3)—supporting the view that the Mec1 pathway mediates DSB interference in *trans* [29]. The more extreme phenotype of *mec1-mn* is likely explained by the fact that Rad24 is not the only activator of Mec1 [7,59–61].

**Figure 7.**
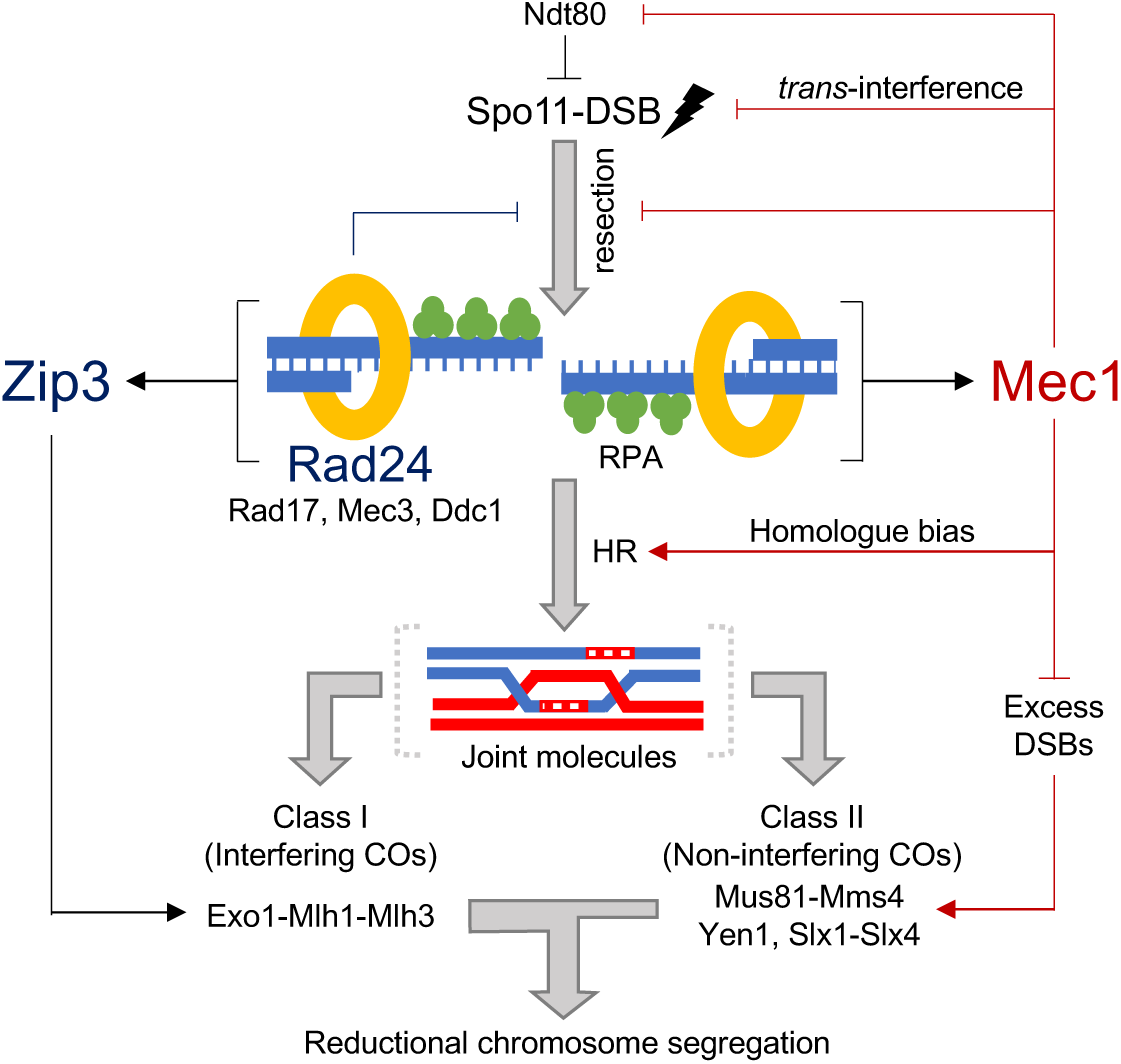
Model of Mec1 and Rad24 activity influencing meiotic DSB formation and repair. Proposed effects of Rad24 are indicated via dark blue lines, and those of Mec1 via dark red lines. Our results support and contribute to the view that Mec1 influences meiotic recombination in numerous ways: promotion of DSB formation via transient inactivation of Ndt80, suppression of DSB formation in *trans*, inhibition of hyper-resection, promoting homologue repair, and suppression of class II CO formation. Rad24 may support these roles by promoting Mec1 activation. In addition, Rad24 promotes Zip3-dependent class I CO formation, and/or the process of CO interference propagation, thereby driving the even distribution of COs along and between chromosomes via the process of CO interference. Together these pathways ensure accurate reductional chromosome segregation during meiosis.

However, in many other cases the phenotypes of *mec1-mn* and *rad24*Δ are less similar, indicating separable roles for the two proteins. For example, the *rad24*Δ mutant is more dependent upon extension of prophase length to maintain high levels of recombination (Fig 2B-D) and spore viability (Fig 1C) than *mec1-mn*. In general, recombination events were also longer upon *RAD24* deletion than in the *mec1-mn* strains (Fig 4D-G), perhaps in part due to small differences in the extent of hyper-resection observed in these two strains [14]. Indeed, Rad24 may enact its role in regulating resection distance by loading the 9-1-1 clamp, which has been proposed to act as a physical barrier to resection (K. Wardell & M. Neale, unpub. obs.). Alternatively or additionally, the effect on resection may be because Rad24 and Rad17 are involved in activating Mec1, which may then prevent resection by deactivation of Sae2 [62] or Exo1 [63]. In vegetative cells, the 9-1-1 complex inhibits resection by promoting the recruitment of Rad9 near DSBs [64]; however, this may not be the case for meiotic DSBs, because Rad9 is not required for checkpoint activation in meiosis [10].

Event length differences may also arise from an alteration in recombination mechanism. Notably, our primary finding is that *rad24*Δ, but not *mec1-mn*, displays a severe defect in the formation of class I COs (Fig 6, discussed below), which is likely a major cause of the low spore viability due to frequent mis-segregation of nonexchange chromosomes. Indeed, in the meioses analysed here, *rad24*Δ strains displayed some of the highest frequencies of non-exchange chromosomes (Fig 2E)—values that are likely to be underestimates of the true non-exchange chromosome frequency due to our selection for meioses generating four viable spores. Such errors are particularly pronounced in the *rad24*Δ *msh2*Δ and *rad24*Δ *sml1*Δ strains, presumably because the lack of checkpoint activation leads to less time to initiate recombination in combination with a class I CO defect. Despite a similar checkpoint defect, we infer that *mec1-mn* strains survive more readily due to a derepression of DSB formation, and no obvious defect in class I CO formation. As indicated above, we recognise that a potential limitation of whole genome recombination assays is the requirement for meioses generating four viable spores, leading to potential selection bias within analysed data. Nevertheless, it seems logical that such biases will tend towards less extreme, more wild type-like phenotypes, suggesting that the characteristics we report here for *rad24*Δ and *mec1-mn* are likely to underestimate the true extent of the defects occurring during meiosis.

### Mixture modelling reveals class I and class II CO events

The methods employed here permit the mathematical separation of class I and class II COs from an otherwise indistinguishable mixture (Fig 6A-F; [37]). Notably, despite similar reductions in the global strength of interference in *rad24*Δ and *mec1-mn* strains, our results indicate that this arises via two distinct mechanisms: excessive class II CO formation in *mec1-mn* strains most likely due to loss of *trans* DSB interference [27,29,65], and what appears to be reduced class I CO formation in *rad24*Δ, most likely due to a defect in Zip3 activation [34] (see below). Importantly, the apparent strength of CO interference between the residual interference-sensitive CO fraction is unchanged in these strains, suggesting that the underlying mechanism that propagates CO interference along chromosomes is not altered (Table S2). It is also interesting to consider that the normal fraction of COs may remain genetically “class I” (i.e. ZMM dependent) in *rad24*Δ, but that sensitivity to CO interference is somehow dysfunctional. Consistent with this alternative hypothesis, the anti-recombinogenic activity of Msh2—which disproportionately targets interfering “class I” COs, skewing their formation to within regions of lower sequence divergence [37]—is still observed within *rad24*Δ (*MSH2* wild type) backgrounds (Fig S11A), suggesting a retention of class I-like identity. By contrast, unlike *rad24*Δ *ndt80AR*, the ability of Msh2 to skew CO position is still present within *mec1-mn ndt80AR* strains—albeit significantly weakened as expected for a mutant with a much higher fraction of insensitive “class II” COs (Fig S11B). Looking forward, we suggest that our mixture modelling techniques may be of great use to assess in more detail the mixed composition of meiotic crossovers present in both published and future datasets of this type, potentially enabling the identification of those components that directly influence the CO interference mechanism.

### Comparisons between the loss of Rad24 and Zip3 function

Loading of Zip3 to chromosomes is partially dependent upon Rad24, but not Mec1 [34]. It may therefore be reasonable to assume that the meiotic defects seen within *rad24*Δ result from a loss of Zip3 efficacy. Consistent with this idea, we demonstrate a significant reduction in the strength of CO interference in *rad24*Δ, but not *mec1-mn* strains (Fig 6). Nevertheless, *rad24*Δ and *zip3*Δ mutants do not appear to present with identical phenotypes, likely due to the pleiotropic nature of the *rad24*Δ mutation that impedes not just class I CO formation via Zip3, but also activation of Mec1 and the meiotic prophase checkpoint.

Direct comparisons made in the *msh2*Δ background reveal (like in *MSH2* strains [48]) that *ZIP3* deletion causes a very significant increase in the event length associated with meiotic COs (Fig S12A)—something that is, in relative terms, increased only a little in *rad24*Δ and *mec1-mn* mutants (Fig S12A). A comparable difference in event length is not observed when considering NCOs (Fig S12B). Larger CO lengths within *zip3*Δ have been proposed to arise from an increase in Sgs1-dependent D-loop extension [48], suggesting that either the residual Zip3 that loads in *rad24*Δ is sufficient to inhibit Sgs1 but is insufficient to mediate class I CO formation, or that Rad24 is required for the spurious Sgs1 activity that arises when Zip3 is completely absent.

*zip3*Δ *msh2*Δ strains also displayed a dramatic reduction in the ratio of COs to NCOs (Fig S12C-D), indicating a loss of CO homeostasis. Whilst *rad24*Δ *msh2*Δ and *rad24*Δ *msh2*Δ *sml1*Δ strains also showed disproportionately few COs, the effect was less severe than observed in *zip3*Δ *msh2*Δ, and, importantly, the paucity of COs in *rad24*Δ was largely restored by extending the length of meiotic prophase (Fig 2C). Taken together, these results suggest a much more significant defect in CO homeostasis upon loss of Zip3 function than upon loss of Rad24.

The overall frequency of recombination is also significantly elevated in *zip3*Δ *msh2*Δ compared to *msh2*Δ (∼206 vs ∼404 events; ∼2-fold; Fig S12C), similar to that observed in *mec1-mn msh2*Δ (∼368 events). Homologue engagement (dependent upon ZMM components) is one of the triggers that suppresses DSB formation at later stages of prophase [66], but the mechanism remains unclear. The similarity between the phenotypes of *zip3*Δ and *mec1-mn*, suggest that homologue engagement may elicit its negative regulatory effect via Mec1. In contrast, despite reduced levels of Zip3 loading within *rad24*Δ [34], no elevation in total event count is observed upon inactivation of this DDR factor (∼188 events; Fig S12C)—suggesting that homologue engagement is still sufficiently active in the absence of Rad24.

Previously in *zip3*Δ mutants, an increase in NCO events formed from dHJ resolution was noted [48], proposed to result from unbiased cutting of dHJ intermediates in the absence of a pathway promoting CO formation. Here we recapture this result in *zip3*Δ *msh2*Δ; the strains display a large increase in NCOs formed from dHJ resolution (Fig S12E). However, overall, only ∼9% of NCOs in *zip3*Δ *msh2*Δ were formed from dHJ resolution, which was actually a significant reduction compared to *msh2*Δ (P<0.05) (Fig S12F). This may be due to an increase in DSB formation due to failure to establish homolog engagement, producing many DSBs that are repaired by SDSA. While there was also a significant increase in the absolute number of NCOs formed from dHJ resolution in *rad24*Δ *msh2*Δ, it was not as significant as the increase in *zip3*Δ or even *mec1-mn* (Fig S12E). However, these NCOs formed a much larger proportion of total NCO counts (∼19%) (Fig S12F). Taken together, these results indicate that in both *zip3*Δ and *rad24*Δ backgrounds, there is an increase in the formation of NCO events from dHJ resolution.

### Meiotic roles of Rad24 and Mec1 orthologues in other organisms

Our findings indicate that Rad24 and Mec1 have both overlapping and distinct functions during *S. cerevisiae* meiosis. Current observations in other organisms suggest relatively conserved roles for ATR^Mec1^ during meiosis (Introduction)—consistent with the coordination between meiotic DNA repair and meiotic chromosome segregation that is essential for fertility. However, there has been relatively little analysis of the roles of the checkpoint clamp and loader during meiosis outside of *S. cerevisiae*.

In *Schizosaccharomyces pombe*, Rad17 (the clamp loader) is required to delay meiotic onset in response to DNA damage, and is important to maintain normal levels of recombination in meiosis and spore viability [67]. Interestingly, *S. pombe* has no Zip3 orthologue, and all COs are non-interfering (class II), being resolved by Mus81 [68,69], suggesting that these defects cannot be mediated via a modulation of the CO interference pathway. In male mice, *Rad9a* mutants display abnormal testes with low sperm counts due to spermatocytes arresting in late zygotene or early pachytene [70]. These defects correlate with a deficiency in DSB repair leading to apoptosis, perhaps due to failure to initiate HR or because of a failure to activate the DDR checkpoint [70]. Similarly, *Hus1* mutant mice display severely reduced fertility and chromosomal abnormalities, but HUS1 is not required for meiotic functions of ATR in response to chromosome synapsis defects [71]. *Hus1* mutant female flies and worms are sterile [72,73], and *Drosophila* Hus1 is required for proper SC disassembly, and efficient DSB processing by HR [74]. In plants, both RAD1 and HUS1 promote accurate DSB repair during meiosis, in particular suppressing nonhomologous end-joining [75], and ZMM-independent CO formation between heterologues [76].

Due to the interplay between various aspects of meiotic recombination regulation, it is somewhat difficult to ascertain which, if any, of these defects are distinct from a generalised abrogation of ATR function. By contrast, building on the work of Shinohara et al [34], our results suggest a critical role for Rad24 during the regulation of CO interference in *S. cerevisiae* (see above)—a function that is clearly specialised for the meiotic system. Moreover, given the essential role that COs play in chromosome segregation and fertility in most sexually reproducing organisms, it will be of great interest to clarify whether or not the DDR checkpoint clamp and/or loader play similar roles across other species as they do in *S. cerevisiae*.

## Acknowledgements

We thank Michael Lichten for strains and methodological suggestions, and Scott Keeney for useful discussions regarding DSB homeostasis.

## Contributions

M.R.C. and M.J.N. conceived the project. M.R.C. performed genome-wide mapping, data processing, event calling, data analysis and interpretation. T.J.C. performed in-silico simulations, mixture modelling and associated data analysis. M.M.K. and B.L. provided scripts, protocols, additional samples, analytical assistance and ideas. M.R.C, T.J.C. and M.J.N. wrote the manuscript.

## Corresponding author

Correspondence to Matthew J. Neale.

## Materials and Methods

### Yeast strains and culture methods

*Saccharomyces cerevisiae* strains used in this study are derivatives of SK1 [77], S288c [78], or BY4741, a derivative of S288c [79]. Hybrid diploid strains were generated by mating haploid SK1 with S288c or BY4741 parents. Strain genotypes are listed in Tables S4 and S5. Gene disruptions were generated by standard Lithium acetate transformation [80]. Gene disruptions of *rad24*Δ::*HphMX, msh2*Δ::*kanMX6, mus81*Δ::*kanMX6* and *zip3*Δ::*HphMX* were performed by PCR mediated gene replacement using pFA6a-*kanMX6* or pFA6-*hphMX* plasmids [81,82]. *P*_*CLB2*_-*MEC1* strains (‘*mec1-mn*’) were created by replacing the natural *MEC1* promoter with the mitosis-specific *CLB2* promoter using pFA6a-*natMX4*-*PCLB2*-*3HA* plasmid as a template [14]. The *P*_*GAL*_-*NDT80*::*TRP1* allele has the natural *NDT80* promoter replaced by the *GAL1*-*10* promoter, and strains include a *GAL4*::*ER* chimeric transactivator for β-oestradiol-induced expression [83]. The base *mlh3*Δ strain (*MATa ura3 lys2 ho*::*LYS2 arg4*Δ(*eco47III-hpa1*) *mlh3*Δ::*kanMX6*) was kindly provided by Michael Lichten.

### Meiotic timecourse with *NDT80* prophase arrest

Diploid strains were incubated at 30 °C on YPD plates for 2 days. For SK1 diploids, a single colony was inoculated into 4 ml YPD (1% yeast extract / 2% peptone / 2% glucose) and incubated at 30°C (250 rpm) for 24 hours. For hybrid crosses, haploid parents are mated in 1 ml YPD for 8 hours, after which an additional 3 ml YPD is added and the cells are grown for a further 16 hours. Cells were inoculated into 30 ml YPA (YPA (1% yeast extract / 2% peptone / 1% K-acetate) to a final density of OD600 = 0.2, and incubated at 30°C (250 rpm) for 14 hours. Cells were collected by centrifugation, washed once in water, then resuspended in 30 ml pre-warmed sporulation media (2% potassium acetate, 5 µg/ml Adenine, 5 µg/ml Arginine, 5 µg/ml Histidine, 15 µg/ml Leucine, 5 µg/ml Tryptophan, 5 µg/ml Uracil), and incubated at 30°C (250 rpm). As necessary, synchronized cultures were split after the required amount of time (e.g. 8 h) in 2% potassium acetate, and one fraction induced to sporulate by addition of beta-oestradiol to a final concentration of 2 mM. Cultures were then incubated to a total of 48 h at 30 °C prior to tetrad dissection.

### Dissection of tetrads/octads to assay spore viability and for sequencing hybrids

To assay spore viability, 50 μl of sporulated cells in sporulation media were incubated with Zymolyase 100T (10 mM Sucrose, 0.7% Glucose, 1 mM HEPES pH 7.5, 1 mg Zymolyase 100T) at a final concentration of 4 µg/ml in a 150 mM sodium phosphate buffer at 37 °C for 15 min. 15 µl of digested cells was pipetted onto a YPD plate and allowed to dry before tetrad dissection. Dissected spores were incubated for 2 days at 30°C on YPD and scored for percentage viability per strain and viable spores per tetrad. To produce hybrid spores for genome sequencing, SK1xS288c and SK1xBY4741 haploid parents were mated for 8–14 hours on YPD plates, with the exception of *ndt80AR* strains, which are mated and grown in liquid YPD for 24 hours (see previous section). Haploids were mated freshly on each occasion and not propagated as diploids, in order to reduce the chance of mitotic recombination. Cells were washed and incubated in sporulation media at 30°C with shaking, and tetrads were dissected after 72 hours. To generate octads, dissected spores were allowed to grow for 4-8 hours on YPD plates until they had completed the first post-meiotic division, after which mother and daughter cells were separated by microdissection, and allowed to grow for a further 48 hours (Figure 1A). Spore clones were subsequently grown for 16 hours in liquid YPD prior to DNA isolation. Only tetrads generating four viable spores, and octads generating eight viable progeny, were used for genotyping by Next Generation Sequencing. The haploid strains used for octad or tetrad sequencing are highlighted in Table S3.

### Preparation of samples for sequencing

To determine the meiotic strand transfers that occurred in the absence of Msh2, we systematically generated and genotyped all members of each octad from S288C x SK1 hybrids. Genomic DNA was purified from YPD cultures of each haploid postmeiotic cell using standard phenol-chloroform extraction techniques. Genomic DNA was diluted to between 0.2-0.3 ng/µl, and concentration measured using the Qubit High Sensitivity dsDNA Assay. Genomic DNA was fragmented, indexed and amplified via the Nextera XT DNA library Prep Kit reference guide workflow according to the Best Practises recommended by Illumina. To check the fragment length distribution and concentration of the purified libraries, 1 μl of undiluted library was analysed on an Agilent Technology 2100 Bioanalyzer using a High Sensitivity DNA chip. Samples were pooled (16-24 per run), denatured by heating at 96°C for 2 minutes, ice-chilled and mixed with denatured PhiX control DNA at 1% final concentration. Sequencing was performed in-house using Illumina MiSeq instruments using paired end 2 × 300 bp or 2 × 75 bp cartridges.

Tetrad analysis was carried out on *ndt80AR* and WT strains, while octad analysis was performed for *msh2Δ* strains. Notably, spore viability in *mec1-mn msh2Δ* and *rad24Δ msh2Δ* backgrounds was still too low for post-meiotic separation to be efficient. Instead, heteroduplex DNA information in *msh2Δ* tetrads was partially retained by harvesting and processing entire colonies as a single sample and categorising variants present in a roughly 50:50 ratio (Figure 5A). This approach allowed the reconstruction of a ‘mocktad’ containing almost all recombination event information that would be present in true octad analysis, with the notable exception of DNA strand orientation. In addition, one *rad24Δ msh2Δ* and one *mec1-mn msh2Δ* tetrad were each restreaked into single colonies and a single colony derived from each of the spores resequenced (Figure 5A). This strategy enabled the full recovery of DNA strand orientation information for these samples (Michael Lichten, pers. comm.).

### Alignment of paired-end reads, detection of SNPs and indels and creation of SK1 genome

Spores were sequenced to an average sequencing depth of 45x. Paired read files are aligned using bowtie2 [84] to both the S288c reference genome (version R64-2-1_20150113) and a custom SK1 reference genome described below. Also included in the reference file are sequences from the yeast mitochondrial (GenBank: KP263414.1 [85]) and 2µ plasmid (GenBank: V01323.1 [86]). To optimize the alignment for long reads and tolerance of mismatches expected in the hybrid genome, the bowtie2 alignment is performed with the following settings: -X 1000 --local --mp 5,1 -D 20 -R 3 -N 1 -L 20 -i S, 1, 0.50. To create a custom SK1 genome, SNP and indel polymorphisms were detected in the S288c alignments using the GenomeAnalysisToolkit (GATK) function HaplotypeCaller [87]. The in-house program ‘VariantCaller’ then combined the GATK calls from 120 samples, in order to calculate the call frequency, total read depth and averaged variant read-depth:total read-depth ratios for each variant. Variants were filtered for a call-frequency between 45-55% of spores, a total read-depth spanning the site of >250 and where 95% or more of the reads at that site contained the variant. Variants located in repeated regions, long terminal repeats, retrotransposons and telomeres were discarded. The final filtered list yielded 64,581 SNPs and 3946 indels. Consistent with polymorphisms being randomly distributed across the *S. cerevisiae* genome, inter-marker distances approximated to an exponential distribution with a mean and median of 169 bp and 81 bp (93.12% of inter-variant distances are <500bp), respectively. The final variant list was then used to automatically produce a custom ‘SK1mod’ genome, using the S288c genome as a backbone and converting any SNP or indel positions into the newly-detected SK1 equivalent. Because this method utilises the S288c reference as a scaffold, called SK1 variants are limited to SNPs and short indels, and therefore lacks any larger structural rearrangements or large insertions/deletions that may exist between the strains.

### Genotype calling in tetrads and octads

Sequence alignment (SAM) files were converted into a sorted binary (BAM) file using the Samtools view command [88], for downstream processing. The PySamStats module ‘variation’ (https://github.com/alimanfoo/pysamstats) was used to produce a list of the number of reads containing an A, C, T, G, insertion or deletion for each genomic position specified in the S288c or SK1 reference, for each spore clone. These whole genome coverage files were filtered using the SNP/indel list derived above and further processed using custom scripts. Genotypes were assigned according to the following rules:

i. All positions must have a read depth of at least 5. ii) A SNP is called as the SK1 variant if 70% or more of reads at that position match the variant, or as the S288c reference if 90% of reads match the reference. iii) If the variant and reference reads are above 90% of all reads *and* within 70% of each other, the position is called as ‘heteroduplex’. iv) Insertions and deletions are called as having the variant genotype if 30% or more of reads at that position match the variant. This low threshold is used because the alignment of indel sequences is biased towards the reference sequence, which means that they are unlikely to be erroneously called as matching the variant genotype. For an indel to be called as the reference genotype, at least 95% of reads must match the reference sequence, and there must be fewer than two reads matching the variant call. This is because even if there is an indel, there are usually reference reads recorded as well due to the difficulty in alignment, so the presence of more than one variant read reduces confidence in the ability to correctly call the position as reference. For this reason, indel positions also cannot be called confidently as heteroduplex. For octads, each of the eight progeny were processed. For tetrads, the four spores were processed and the data for each spore duplicated in order to appear as an octad in order to simplify downstream processing. In octads and MMR-proficient tetrads, SNPs called as heteroduplex were discarded. However, in MMR-deficient tetrads, heteroduplex calls are converted as described below. To improve the fidelity of this method, SNP positions found to contain mixed reads across multiple (>2) samples were eliminated— presumed to be residual errors in the variant table. For each position with a heteroduplex call, the original ‘mother’ is converted to SK1 and the duplicated ‘daughter’ to S288c. This is an arbitrary choice since it is impossible to know which should have which call, assuming there is no bias in the directionality of the repair. The exceptions are *rad24*Δ *msh2*Δ #6 and *mec-mn msh2*Δ #3 in Table S1, in which a single member of the mother-daughter pair were resequenced in addition to the mixed reads (see above). This latter strategy enables the haplotype of the other member of the mother-daughter pair to be inferred. In all cases, the genotype calls are converted into a binary signal, either 1 for S288c or 0 for SK1.

### Event Calling

Event calling utilized a custom Python script as described below [35,36]. Using the binarized input, chromosomes were split into segments with the same segregation pattern [36]. Also recorded is the segment type, which will be 1:7, 2:6, 2:6*, 3:5, 4:4, 4:4*, 5:3, 6:2, 6:2* or 7:1 as described [35]. Recombination events were called as being a set of segments located between two 4:4 segments longer than 1.5 kb [36]. A 4:4 segment corresponds to a Mendelian segregation profile, 5:3 and 3:5 segments to half-conversion tracts, 6:2 and 2:6 segments to full conversion tracts, etc. Each recombination event can contain between 0-2 COs or NCOs. Additionally, we encounter events which cannot be classified because they occur at the end of the chromosome and never return to a 4:4 pattern – these are given type ‘U’. Figure 2A-D show the total number of COs and NCOs present in the events (counting two COs or NCOs when necessary). The events are also classified by the number of chromatids involved: one chromatid, two chromatids, either sister or non-sister, and three or four chromatids. All recombination event images are provided (Supplementary files). Alongside the recombination event strand patterns, the Spo11 profile and hotspot strength values [89], Rec114/Mer2/Mei1 (RMM) profile [90], Rec8 peaks [91], and transcribed regions [92] are plotted. Chromatid strand orientation in octads was reconstructed as described [36] and where it could be determined, is displayed in event images (**Supplementary Files**). Events were further subclassified according to strand transfer patterns present (Fig S5 and S6) as described [36]). Patterns include restoration patches, full-conversion patches and bidirectional conversions.

### Event tract length determination

The event mid-length, defined as the distance between the mid-points of the first and last markers to be converted at each end of the event, was used as an estimate of the true event tract length (Fig 4A). Multi-DSB events are excluded from the median calculation due to the difficulty in determining how much of the recombination tract length is attributable to each component event.

### Quality control regarding mitotic events

One *mec1-mn ndt80AR* tetrad (#6) contained a number of events that appeared to be mitotic. The tetrad in question contained three large (>100kb) regions of 8-0 segregation on chromosomes 4, 7 and 10 (Fig S13A), as well as several smaller 8-0 regions (event images in **Supplementary File “TCMN6_Annotated_Images.pdf”**). These are considered likely to be premeiotic conversions due to the lack of accompanying hDNA, and the tetrad was thus excluded from most analyses except for the examination of event tract lengths in the rest of the genome. One *mec1-mn msh2*Δ tetrad (#1) also had an apparent premeiotic event, although of a different nature. The presence of heteroduplex reads along the entire length of chromosome 8 in two of the four spores suggests that these two spores contain two copies of chromosome 8, one from each parent, suggesting that a duplication of the S288c copy of the whole chromosome had occurred just prior to entering meiosis (Fig S13B). This tetrad was also excluded from other analyses except for event tract lengths on the unaffected chromosomes.

As 8:0 segregation patterns may arise from rare mitotic recombination events occurring prior to meiosis, 8:0 segments were discarded from analyses if they had no adjacent hDNA, or appeared in the same locus across multiple meioses—suggesting mis-annotation within one of the parental genomic references. Moreover, to limit the possibility of mitotic recombination events contaminating meiotic patterns, haploids were mated freshly for each experiment, rather than propagating the strains as diploids, and mating before sporulation was limited to <14 hours.

### Determining hotspot overlap of one-sided noncrossover events

The maximum possible start and stop coordinates for each one-sided NCO event were used and compared to a list of Spo11-DSB hotspots [89]. If a hotspot intersected with the maximum possible event region, the event was considered to have hotspot overlap.

### Inter-crossover distance distributions and mixture modelling

In order to model/quantitatively describe CO distributions, inter-crossover distances (ICDs) were fit with a gamma (γ) distribution function – a multivariate, continuous probability distribution characterised by two independent parameters: (i) (γ)α (shape factor) (ii) (γ)β (scale factor). A value of (γ)α=1 indicates an exponential distribution i.e. randomness/no interference. (γ)α values of >1 indicate interference. (γ) distributions naturally arise when considering intervals between successive, independent events and thus represent an ideal system to model ICDs. The extent to which a system has deviated from randomness (i.e. is interfering) is typically proportional to the value of (γ)α. Maximum likelihood estimate (MLE) gamma-fitting was performed using the MATLAB (2018a) function *fitdist*.

To obtain simulated ICDs, virtual recombination events were sequentially placed along simulated chromosomes using an in-house script (‘RecombineSim’, MATLAB 2018a) under varying conditions (i) independence, whereby all events are randomly placed (ii) interference, whereby an event imposes a bidirectional interference window (hazard function) derived from best-fit gamma parameters as previously described [93], reducing the probability of a second proximal event. A fractional amount of class II COs that remain insensitive to this window were introduced via the C_PROB_ parameter where necessary. All simulations (N = 10,000 cells), by design, precisely matched the event count experimentally observed for any given genotype/cell. Any simulated events residing within a 1.5 kb window of each other were merged.

Experimental inter-CO distance data is expected to comprise of two subpopulations: (i) a random, class II CO component (γ)α=1 (ii) a non-random, interfering class II CO component (γ)α>1. To statistically estimate/separate these latent variables, an in-house mixture modelling algorithm (‘GMM’), based on expectation maximisation, was utilised to fit two gamma distributions to the overall data (one for each CO subclass). The quality of fit is iteratively and progressively improved as the system converges on the maximum likelihood estimate to provide gamma parameter ((γ)α, (γ)β)) estimates for each subcomponent as well as the relative contribution of each subcomponent to the overall fit (i.e. the estimated class I:class II ratio).

## Statistical analyses

The *Kolmogorov–Smirnov (KS)* test, performed with the MATLAB (2018a) function *kstest/kstest2*, is a nonparametric test used to compare continuous probability distributions. A KS-test may therefore be used to compare inter-event distance cumulative frequency distributions either between empirical samples (two-sample test) or to compare a sample to a hypothetical gamma distribution (one-sample test). *Fisher’s exact test* was performed with the R environment (http://www.Rproject.org/) using the R function fisher.test(),with the two-sided option. *Wilcoxon’s test* was performed with the R environment using the R function wilcox.test(), with the two-sided option and a continuity correction. *Student’s T-test* was performed with the R environment using the R function t.test() and the two-sided option.

## Data availability

All strains listed in Tables S2 and S3 are available upon request. Octad and tetrad sequences are publicly available at the NCBI Sequence Read Archive (accession numbers **SRP151982, SRP152540, SRP152953**). Scripts are available on request.

## Supplementary Figures

**Figure S1.**
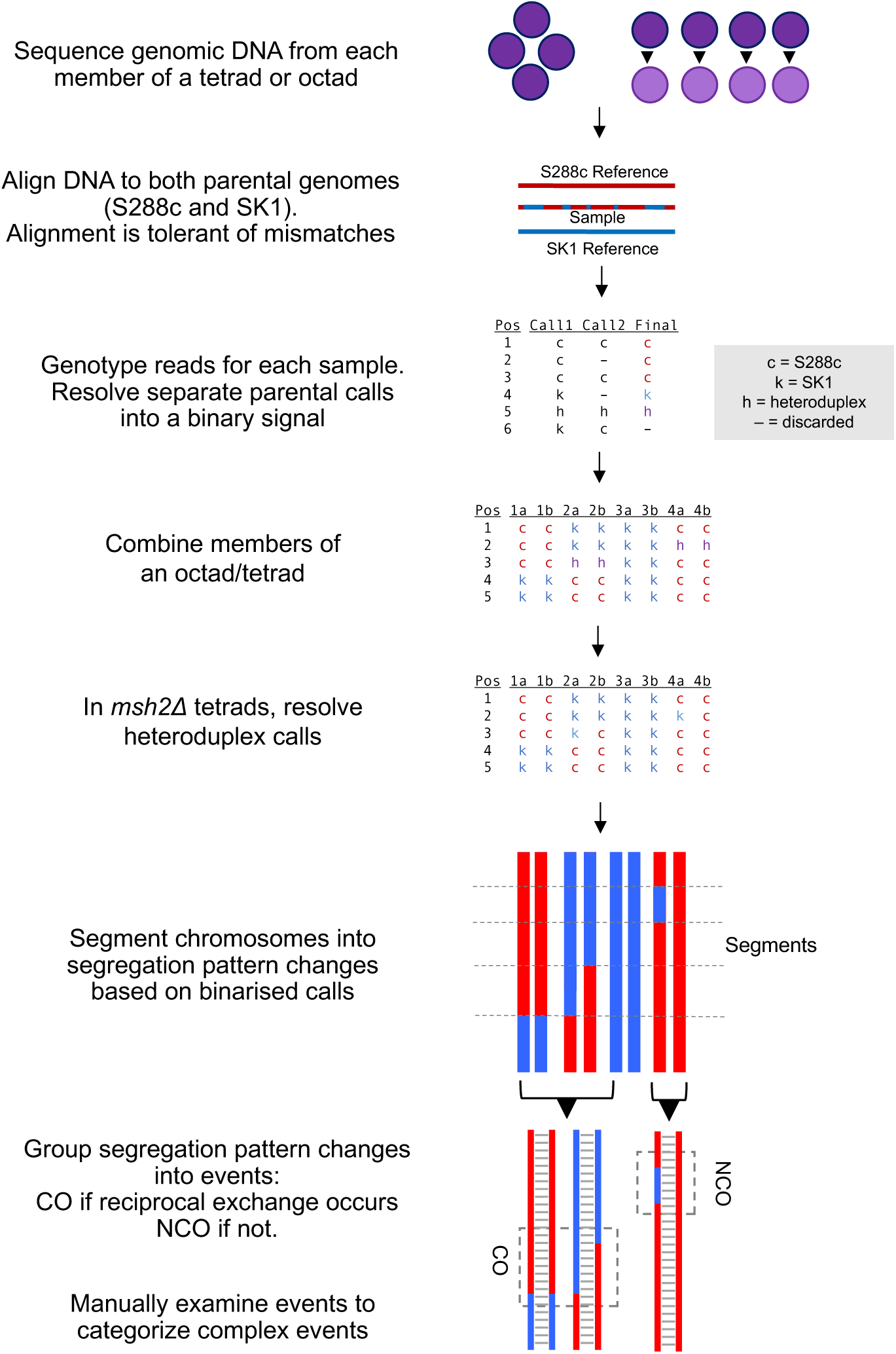
Bioinformatics pipeline for whole-genome mapping of meiotic HR events. The method is based on that described previously [35]. Notable differences are that mismatches are tolerated during the initial alignment to preserve reads containing variants from both parents; the alignment is carried out against both parental genomes, and genotype calls reconciled; and the variant table includes both SNPs and indels. Additionally, we introduce the technique of sequencing *msh2*Δ tetrads and using heteroduplex calls to reconstruct an ersatz octad.

**Figure S2.**
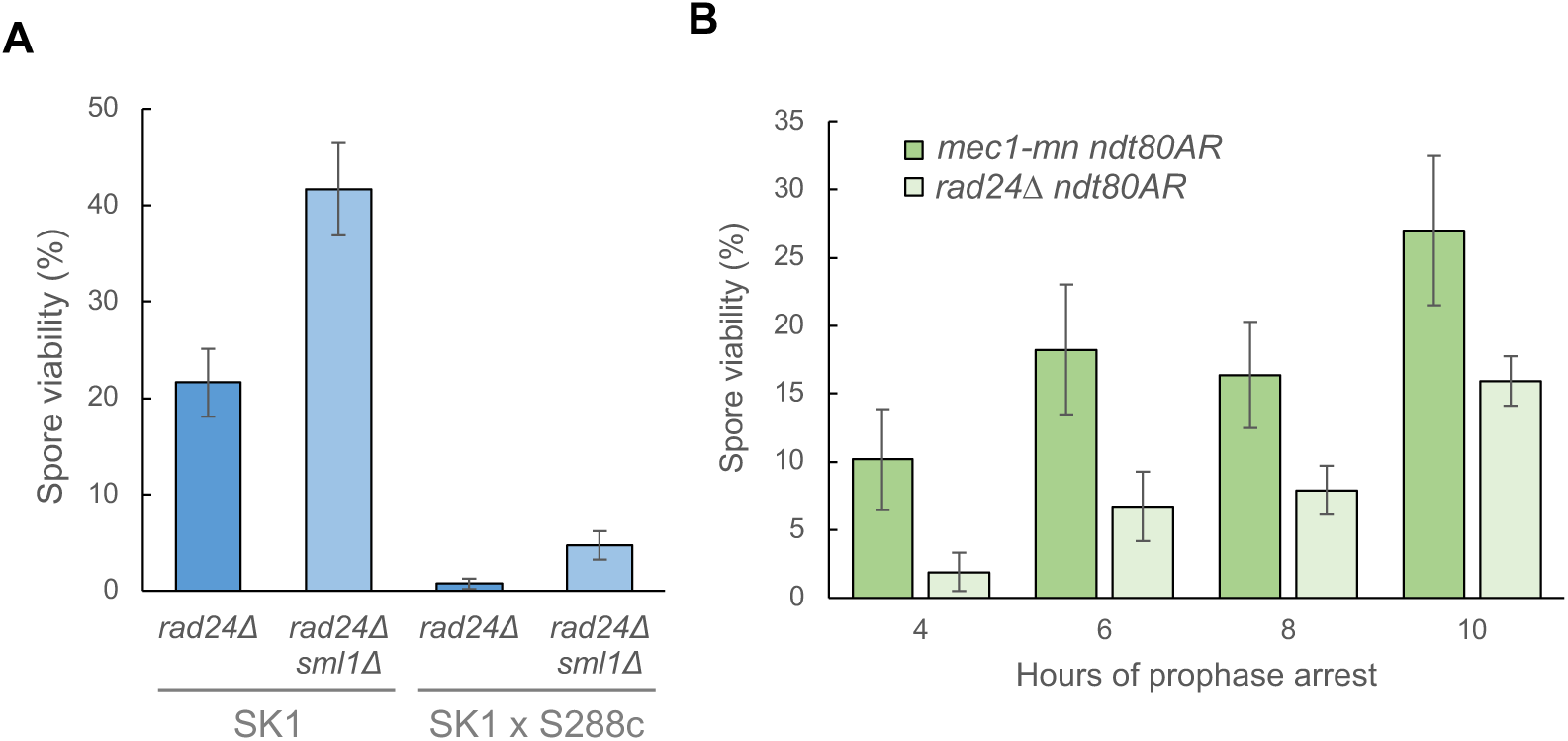
Methods used to increase *rad24*Δ and *mec1-mn* spore viability. All error bars are standard error of the mean. **A)** Rescue of *rad24*Δ spore viability by *sml1*Δ. The spore viability of both SK1 and SK1xS288c hybrid strains with a deletion of *RAD24* is improved by the additional deletion of *SML1*. **B)** The effect of increasing prophase length on the spore viability of hybrid *mec1-mn* and *rad24*Δ yeast. Hybrid SK1xS288c strains are arrested in prophase and released after 4-10 hours using an inducible *NDT80* system ‘ndt80AR’. Both Mec1 and Rad24 mutants display an improvement in spore viability when prophase is extended (compare with non-arrested viabilities shown in Fig 1C), and the level of improvement correlates with the length of the arrest.

**Figure S3.**
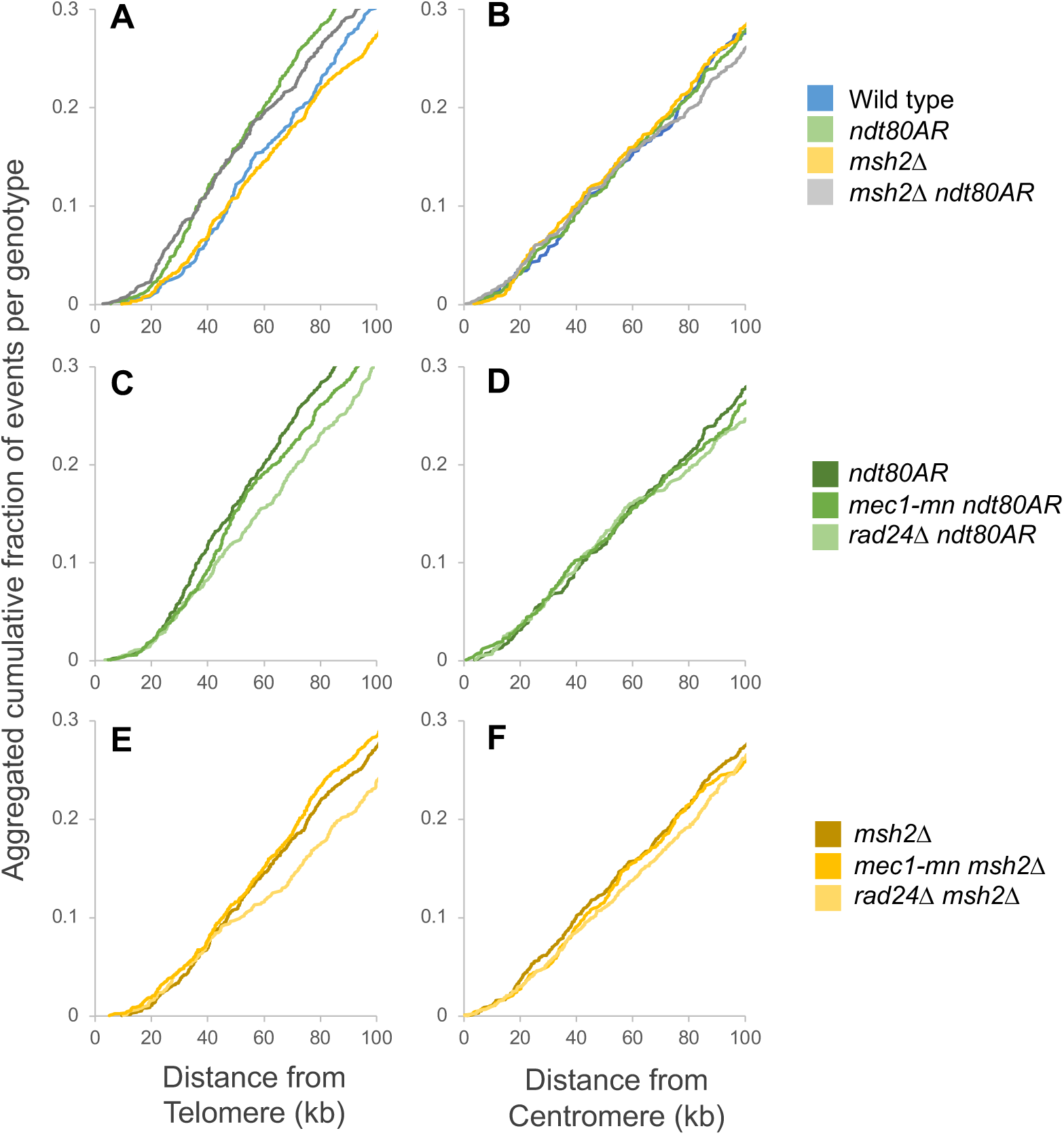
Prophase arrest skews recombination event formation towards telomeric regions. The cumulative fraction of all recombination events plotted against the distance from the nearest telomere **(A,C,E)** or centromere **(B,D,F)**, for the indicated strains.

**Figure S4.**
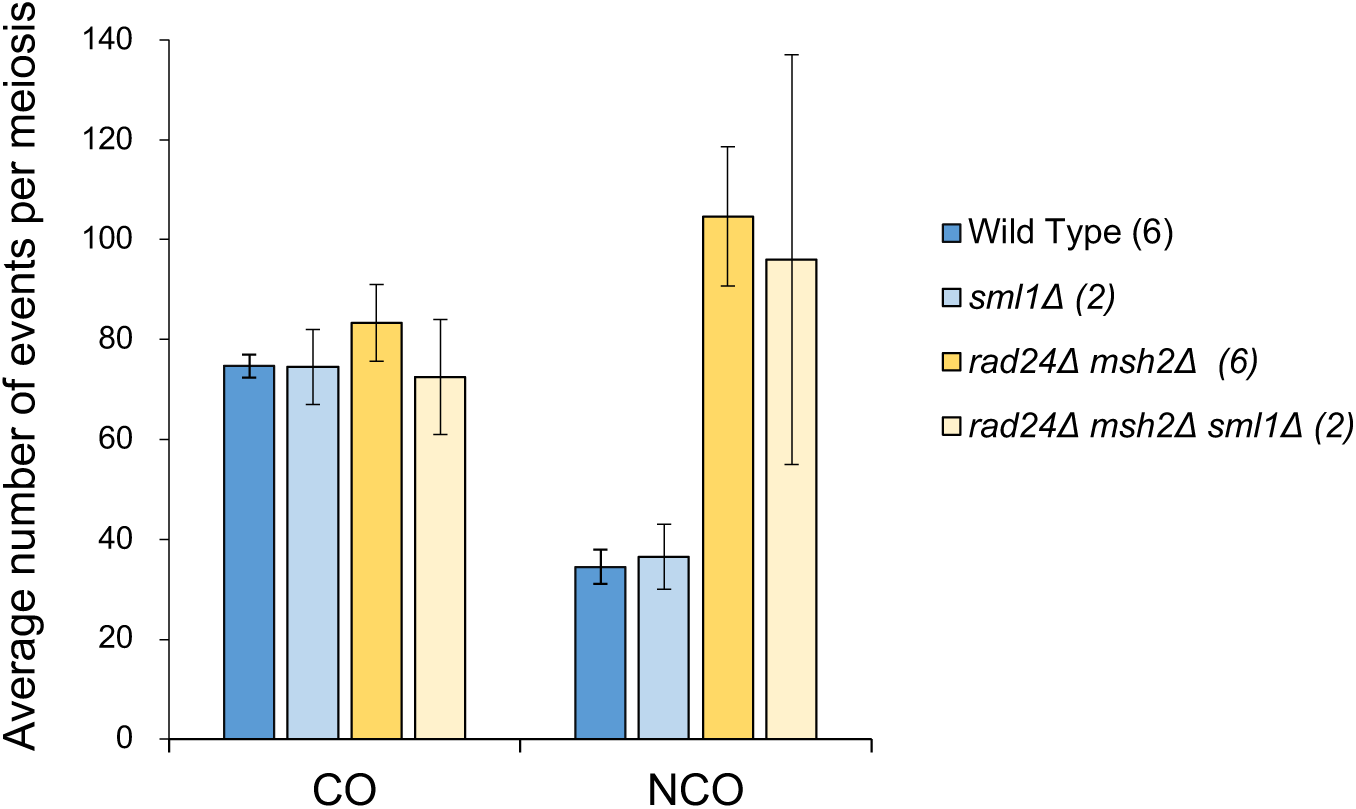
Meiotic recombination frequency is unchanged by *sml1*Δ. The mean counts of CO and NCO events per tetrad are shown (including both single and multi-DSB events). All error bars are standard error of the mean. Event count differences between *sml1*Δ and *SML1* versions of strains were tested by two-tailed T-test, and found to be insignificant.

**Figure S5.**
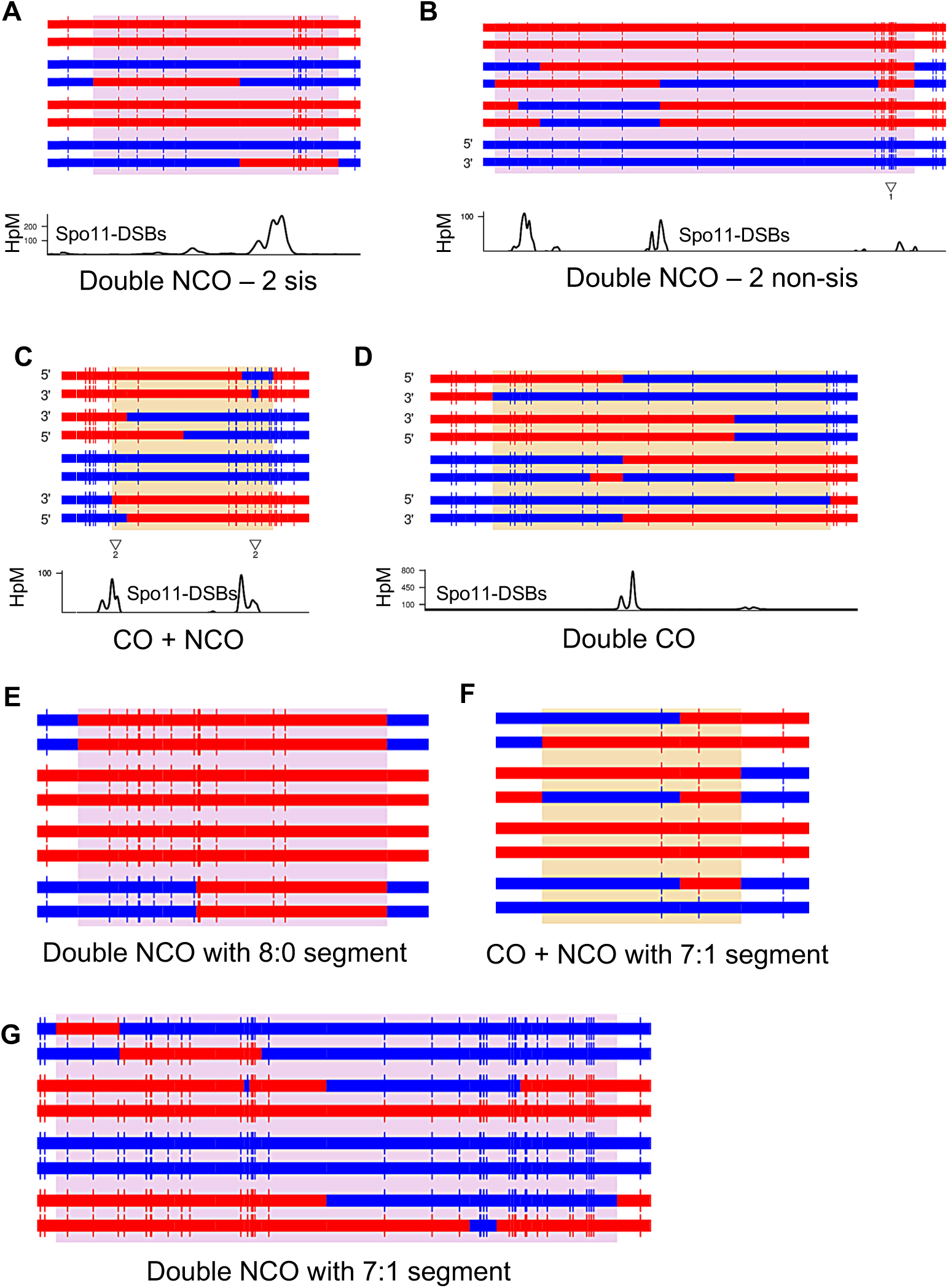
Examples of events matching multi-DSB categories. All example images are from *msh2*Δ octads. **(A-G)** A representative set of multi-DSB events taken from *msh2*Δ octads. Horizontal lines represent the eight strands of DNA present during recombination, while vertical lines are SNP/indel locations, with the S288C and SK1 alleles coloured red and blue, respectively. The orange/purple background highlights the CO/NCO event region respectively; events containing both a CO and an NCO are also coloured orange. The bottom part of panels **A-D** shows the counts of immunoprecipitated Spo11-FLAG oligos for each position, using S288C coordinates [89] smoothed using a 101bp hanning window. **A)** Double noncrossover affecting two sister chromatids. **B)** Double non-crossover affecting two non-sister chromatids; this is determined by the occurrence of a small region of double-exchange, which could potentially be compatible with a very close double CO. **C)** CO and NCO event; this is determined by the occurrence of the NCO on a separate chromatid to the CO event. **D)** Double CO event; this is determined by the double reciprocal exchange. **E)** Double noncrossover affecting two sister chromatids, and containing a region of 8:0 segregation. **F)** CO and NCO event containing a region of 7:1 segregation. **G)** Double NCO containing a region of 7:1 segregation.

**Figure S6.**
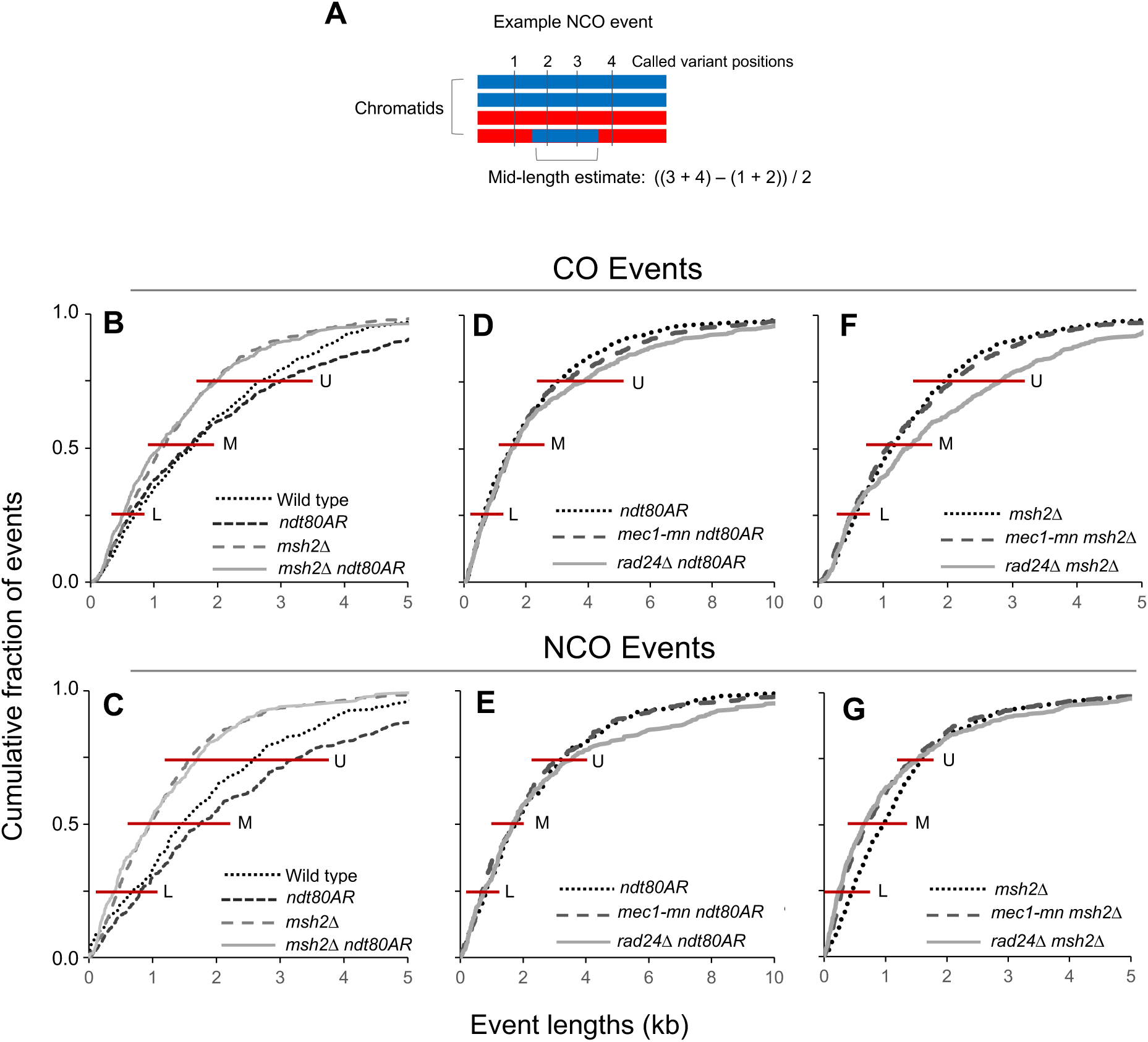
Meiotic recombination event tract lengths in the absence of Mec1 or Rad24. **A)** Event mid-length calculation is defined as the distance between the midpoints of the intermarker intervals surrounding the recombination event. Multi-DSB events are omitted from these analyses. **B-G)** Distribution of mid-lengths of the strand transfers associated with CO **(B,D,F)** and NCO **(C,E,G)** events in the indicated strains. Red horizontal lines indicate the values of the Upper Quartile (U), Median (M) and Lower Quartile (L) summarised in Fig 4.

**Figure S7.**
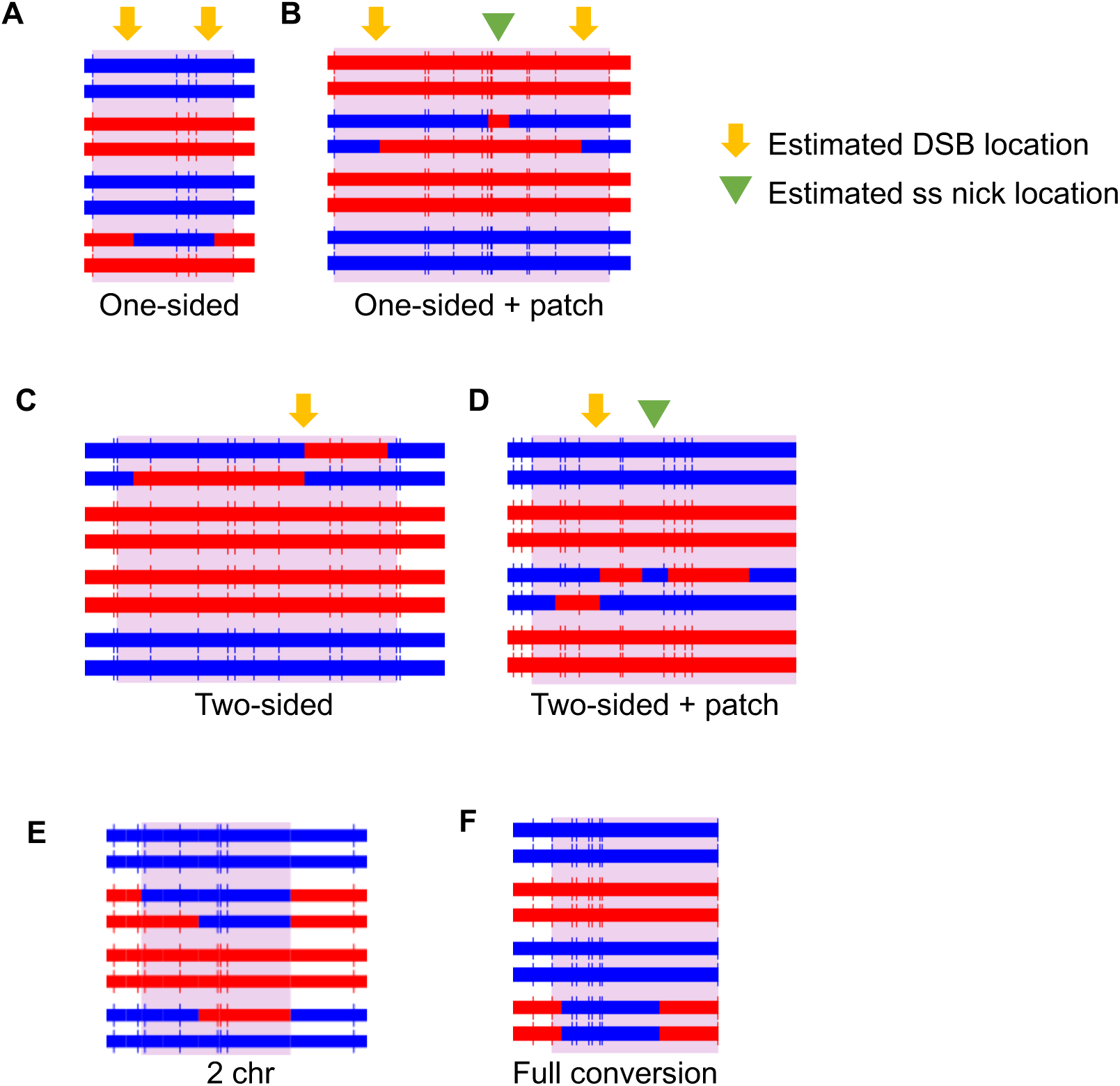
Examples of events matching NCO categories. All example images are from *msh2*Δ octads. **(A-F)** A representative set of NCO events taken from *msh2*Δ octads. Horizontal lines represent the eight strands of DNA present during recombination, while vertical lines are SNP/indel locations, with the S288C and SK1 alleles coloured red and blue, respectively. The purple background highlights the NCO event region. Categories are named as in [36]. The term ‘Two-Sided’ refers occurrence of strand transfer patterns on both sides of the putative DSB location. A lack of hDNA on both sides of the event (‘One-Sided’) may be caused by an absence of markers, or may be due to a peculiarity of the repair process e.g. template switching. **A)** One-sided noncrossover with a half-conversion tract. **B)** One-sided noncrossover with a half-conversion tract and an internal patch of full conversion. **C)** Two-sided noncrossover with two half-conversion tracts (*trans* hDNA) affecting the same chromatid. **D)** Two-sided noncrossover with two half-conversion tracts affecting the same chromatid and separated by a restoration tract. **E)** Noncrossover that affects two non-sister chromatids. **F)** Noncrossover with a single full-conversion tract.

**Figure S8.**
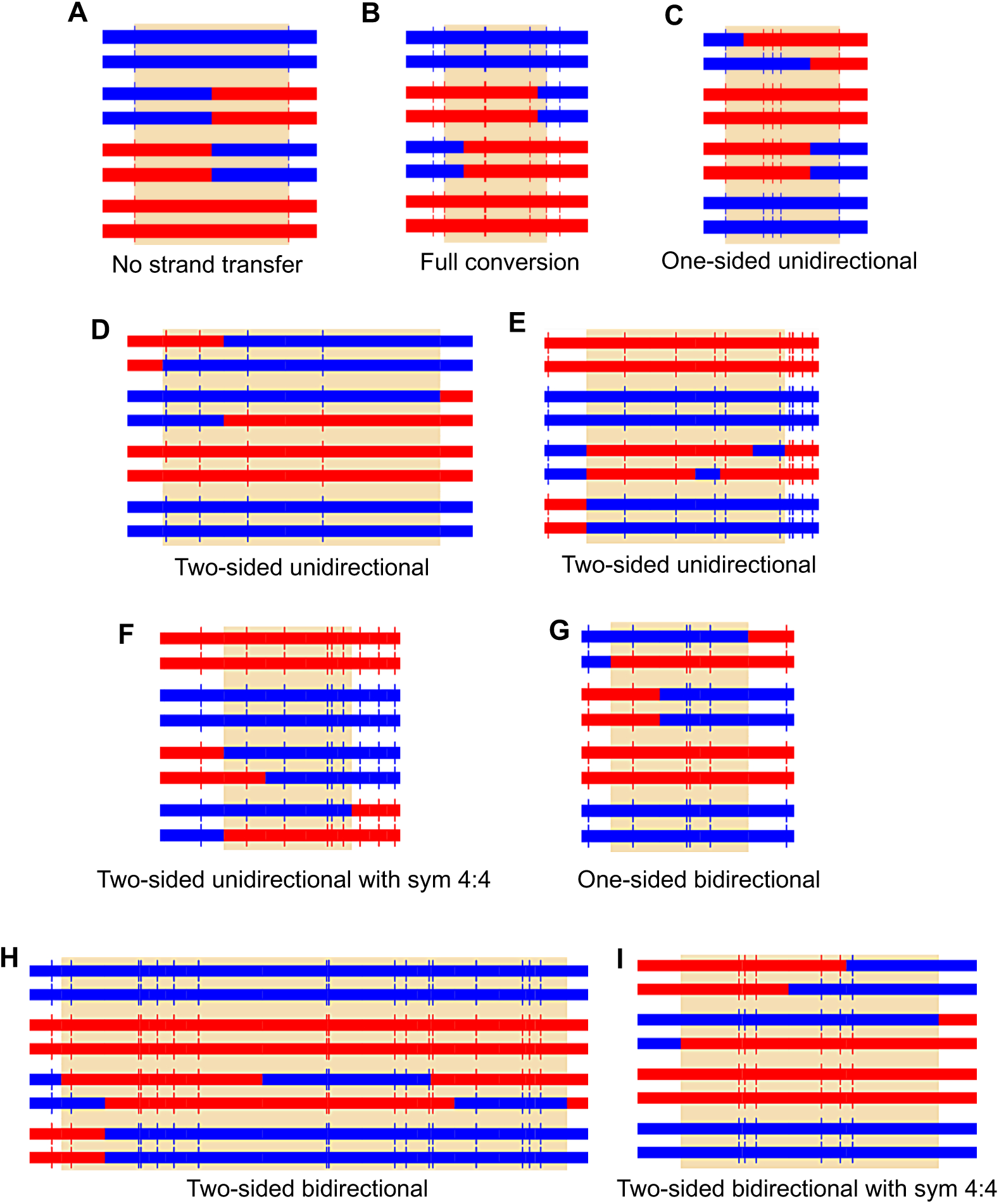
Examples of events matching CO categories. All example images are from *msh2*Δ octads. **(A-I)** A representative set of CO events taken from *msh2*Δ octads. Horizontal lines represent the eight strands of DNA present during recombination, while vertical lines are SNP/indel locations, with the S288C and SK1 alleles coloured red and blue, respectively. The orange background highlights the CO event region. Categories are named as in [36]. The term ‘Two-’ or ‘One-Sided’ refers to the occurrence of strand transfer patterns on one or both sides of the putative DSB location. The term ‘Directionality’ refers to whether markers from only one parent are converted, or both. Bidirectionality may be caused by multiple DSBs/nicking, or junction migration. ‘Sym 4:4’ refers to symmetrical hDNA, which may originate from HJ branch migration. **A)** Crossover with no detectable strand transfer. **B)** Crossover with a single full-conversion tract. **C)** One-sided, unidirectional crossover with a single half-conversion tract. **D)** Two-sided, unidirectional crossover with trans hDNA tracts on the two recombining chromatids. **E)** Two-sided, unidirectional crossover with trans hDNA tracts on one recombining chromatid. **F)** Two-sided, unidirectional crossover, with a tract of symmetrical hDNA. **G)** One-sided, bidirectional crossover. **H)** Two-sided, bidirectional crossover with trans hDNA tracts on one chromatid only. **I)** Two-sided, bidirectional crossover with trans hDNA tracts on both chromatids, and a tract of symmetrical hDNA.

**Figure S9.**
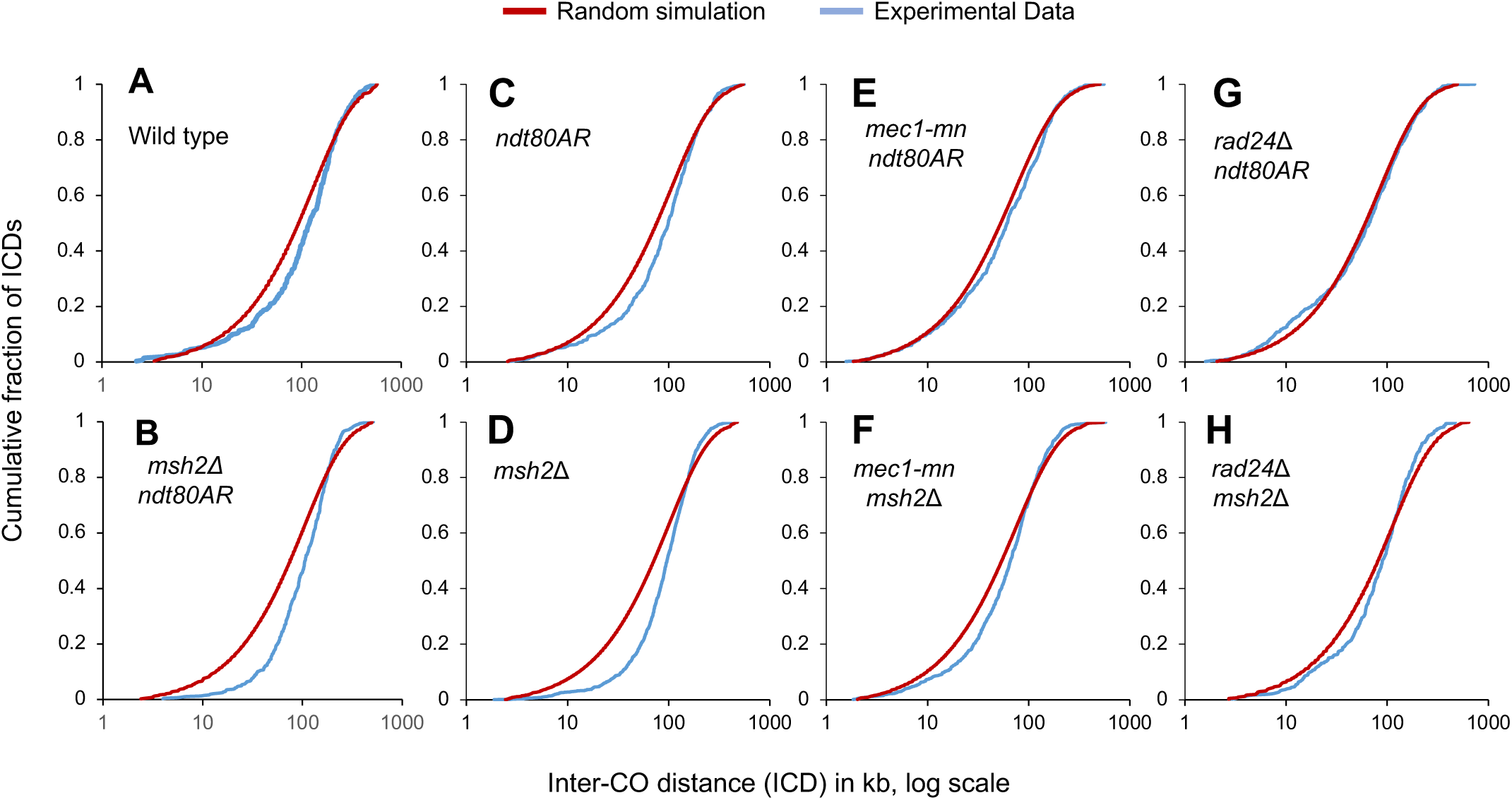
Inter-CO distance is not altered by extending prophase length, but is increased by *msh2*Δ. **A-H)** Inter-CO distances (ICDs) were calculated for the indicated strains, arranged in rank order from smallest to largest and plotted as cumulative fraction of the total ICD count against inter-CO distance on a log scale. For comparison, a simulated dataset was produced where CO events were placed randomly for the same number of CO events as were observed each strain background.

**Figure S10.**
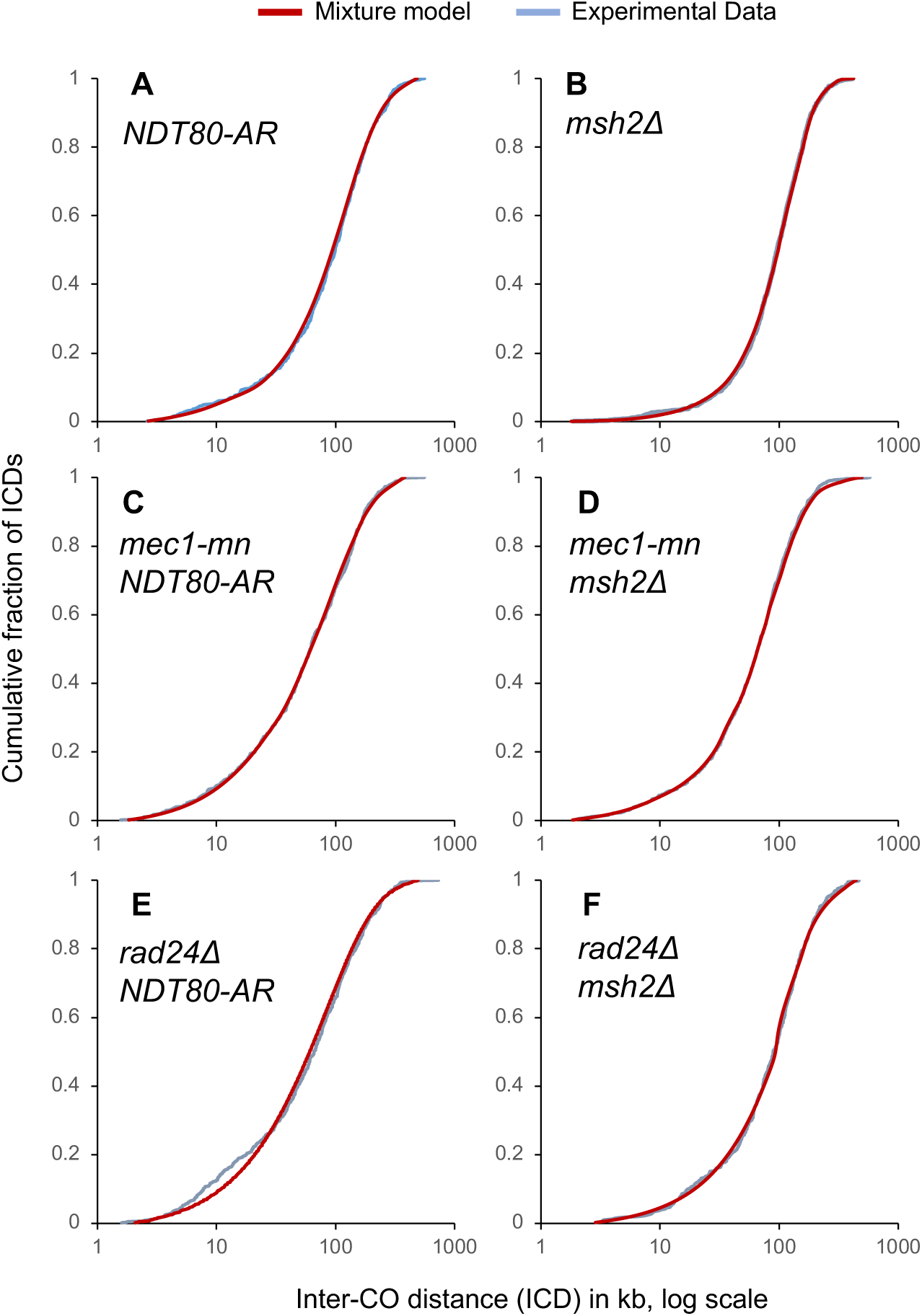
Inter-CO distance distributions can be accurately recaptured by a computational model. Experimentally-derived inter-CO distances (ICDs) were calculated for *mec1-mn* and *rad24Δ* strains in an *ndt80AR* **(A,C,E)** or *msh2*Δ **(B,D,F)** background, arranged in rank order from smallest to largest and plotted as cumulative fraction of total ICD events against ICD size on a log scale. The ICD distribution for each strain was fitted to a mixture model, which extracts a random and an interfering distribution (**Methods**; [37]). A simulated dataset was produced (**Methods**; [37]) where COs were placed under the influence of interference using parameters determined by the mixture model (Table S4), for the same number of CO events as were in each strain background, and plotted alongside the experimental data. This allows the goodness-of-fit of the model to be visually determined. In the *rad24*Δ *ndt80AR* strain, two separate distributions could not be extracted; it appears to have a single, completely random component.

**Figure S11.**
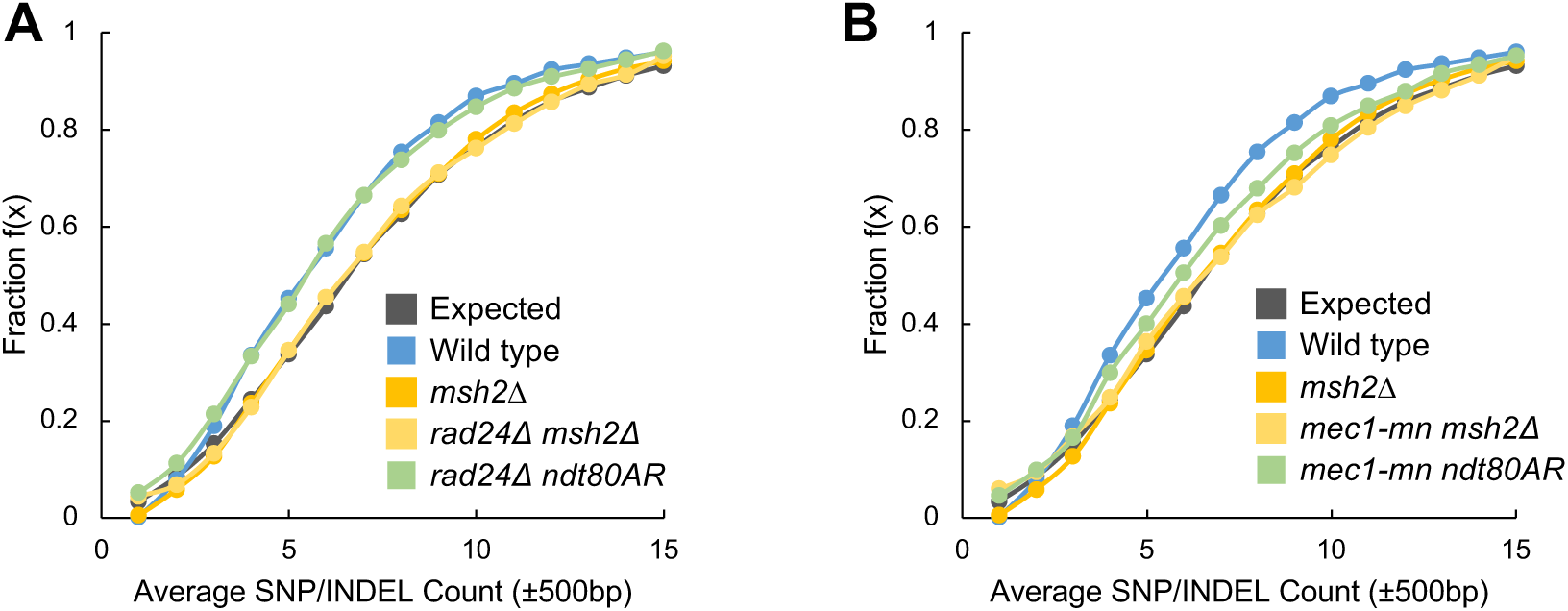
COs within *rad24*Δ retain class I-like identity. **A-B)** Empirical cumulative distribution functions (eCDFs) showing the fraction of COs that reside within a region of a given SNP/INDEL count for each specified genotype (**Methods**; [37]). SNP/INDEL count is assayed using a ±500bp window centered on CO or DSB hotspot midpoints. All contained SNP/INDELs are then subsequently tallied, with equal weight. Expected is calculated using DSB hotspot midpoints [94].

**Figure S12.**
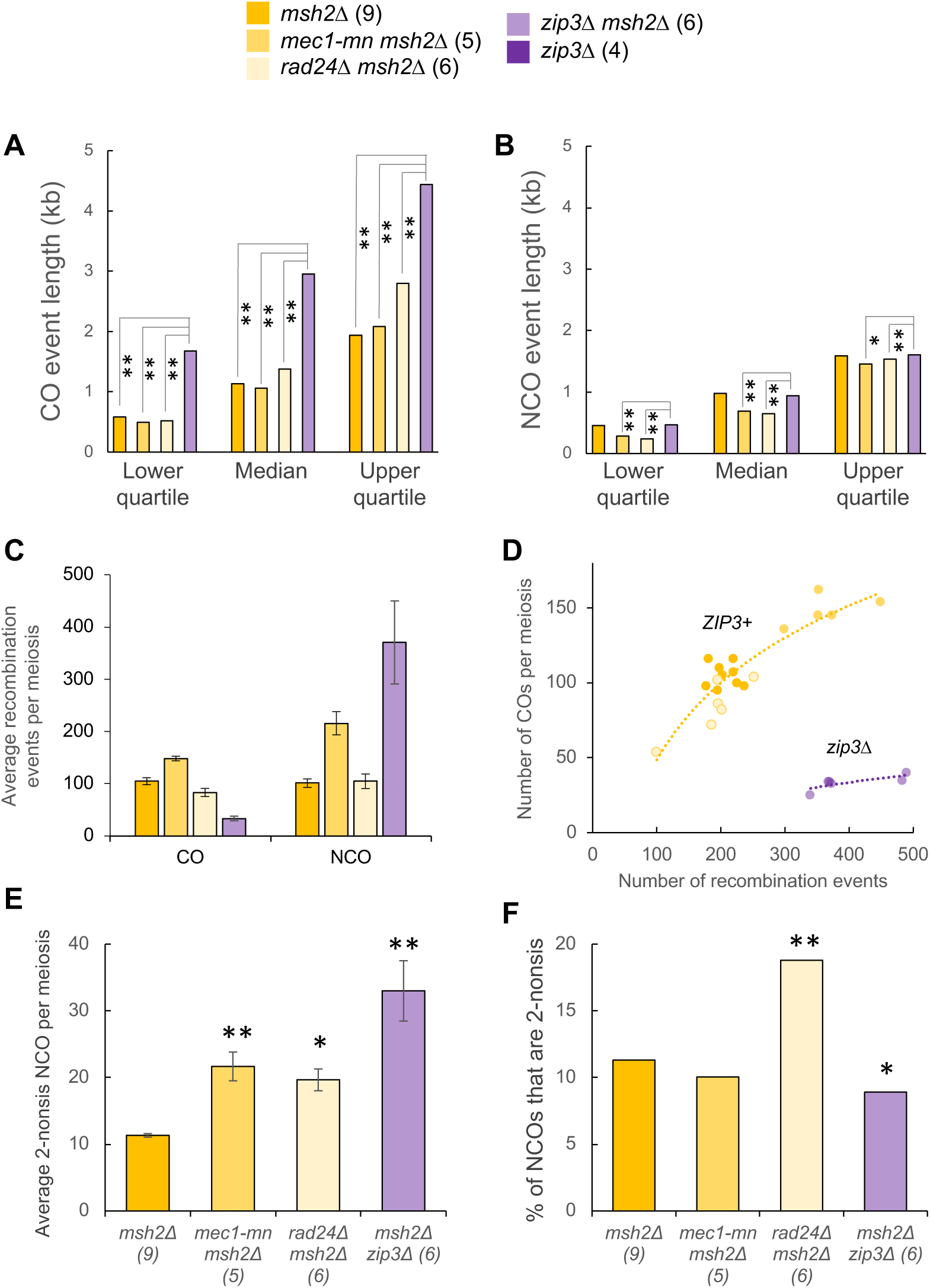
Comparison of recombination event characteristics between DDR mutants and *zip3*Δ. **A)** CO and **B)** NCO event lengths plotted as a cumulative frequency distribution. Multi-DSB event lengths are omitted. **C)** Recombination event counts. **D)** The relationship between recombination event counts and the number of CO events in individual meiosis. The lines are an exponential model for either the *zip3*Δ *msh2*Δ strain (purple) or all other (*ZIP3*+) strains (orange). **E)** The average number of NCO events that appear to have been formed by dHJ resolution. Differences to the reference strain were tested by T-test (*:P<0.05, **:P<0.01). **F)** The aggregate proportion of all NCO events that appear to have been formed by dHJ resolution. Differences to the reference strain were tested with Fisher’s Exact Test (*:P<0.05, **:P<0.01).

**Figure S13.**
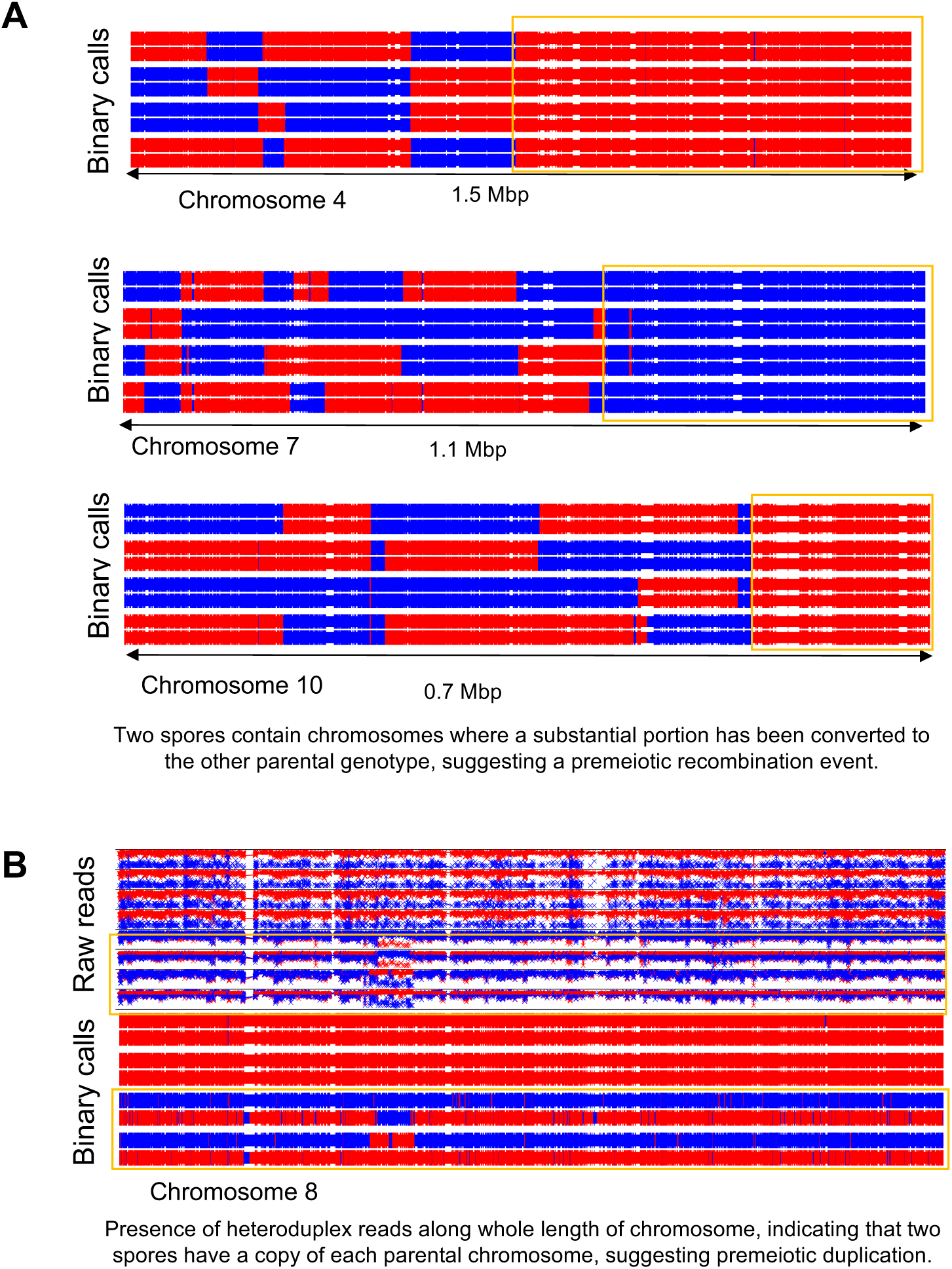
Abnormal mitotic recombination events in *mec1-mn* strains. Whole chromosome recombination patterns were mapped in hybrid strains by detection of SK1 (blue) or S288c (red) markers. The region of interest is plotted as eight horizontal lines corresponding to the eight strands of DNA present in the original hybrid diploid, with vertical tick marks indicating called variant positions. **A)** partial chromosome conversions on Chr 4, 7 and 10 in a *P*_*CLB2*_*-MEC1* +10 h tetrad (TCMN6); **B)** A duplication of S288c Chr 8 in a *P*_*CLB2*_*-MEC1 msh2Δ* tetrad (TCMM4). Raw reads indicate the frequency of reads containing SK1 or S288c type polymorphisms detected at each position; these are translated into binary calls, which can only be SK1 or S288c.

**Table S1.** Key characteristics of meiotic recombination in sequenced meioses. For each individual meiosis, values are given for the number of single CO and NCO events, the identity of chromosomes without a CO or NCO event, the number of multi-DSB events (either dCO, dNCO or CO + NCO), and the number of individual events containing an 8:0 or 7:1 segment (not an exclusive category; all such events are also considered to be multi-DSB events of some kind). An increase in the formation of multi-DSB events suggests a loss of CO interference and/or *cis/trans* DSB interference. The occurrence of chromosomes without COs suggests a loss of CO assurance; COs are not spread out evenly between chromosomes. Many of these chromos are also missing NCOs, meaning they could not have compensated for the lack of CO events. The occurrence of 7:1 or 8:0 segregation patterns within an event suggests a loss of *trans* DSB interference. Example images for multi-DSB and 8:1/7:1 segment events are shown in Figure S4. *\* mec1-mn msh2Δ 1* and *mec1-mn ndt80AR* 6 were not used for any analysis except event lengths, due to large mitotic duplication events (see **Methods**, Fig S13). *rad24*Δ *msh2*Δ #6 and *mec-mn msh2*Δ #3 underwent haplotype resequencing (**Methods**) to reproduce a full octad (see Fig 5A).

| Meiosis | Single COs | Single NCOs | Chromosomes without a CO | Chromosomes without an NCO | Double COs | Double NCOs | CO and NCO | 8:0 segregation | 7:1 segregation |
| --- | --- | --- | --- | --- | --- | --- | --- | --- | --- |
| Wild type 1 | 68 | 35 | Chr 1, 6 | Chr 1, 3, 6 | 0 | 2 | 3 | 1 | 11 |
| Wild type 2 | 83 | 40 | Chr 1 | Chr 6 | 0 | 1 | 3 | 0 | 15 |
| Wild type 3 | 76 | 36 | - | Chr 2, 3 | 0 | 2 | 4 | 1 | 13 |
| Wild type 4 | 71 | 33 | - | Chr 1, 5, 6, 12 | 0 | 2 | 3 | 1 | 12 |
| Wild type 5 | 71 | 29 | Chr 1 | Chr 3, 5, 8 | 0 | 0 | 1 | 0 | 12 |
| Wild type 6 | 77 | 18 | - | Chr 6, 7, 9, 10, 11 | 1 | 1 | 2 | 4 | 8 |
| *msh2Δ* 1 | 116 | 98 | - | - | 0 | 3 | 7 | 0 | 0 |
| *msh2Δ* 2 | 100 | 123 | - | - | 0 | 1 | 6 | 0 | 0 |
| *msh2Δ* 3 | 95 | 88 | - | - | 0 | 6 | 5 | 0 | 0 |
| *msh2Δ* 4 | 103 | 92 | - | - | 1 | 3 | 10 | 0 | 1 |
| *msh2Δ* 5 | 98 | 77 | - | Chr 6 | 0 | 1 | 0 | 0 | 0 |
| *msh2Δ* 6 | 114 | 61 | - | Chr 1, 7 | 1 | 2 | 1 | 1 | 0 |
| *msh2Δ* 7 | 98 | 131 | - | - | 0 | 4 | 10 | 0 | 1 |
| *msh2Δ* 8 | 107 | 105 | - | Chr 6 | 0 | 4 | 9 | 0 | 2 |
| *msh2Δ* 9 | 110 | 86 | - | - | 0 | 1 | 7 | 0 | 0 |
| *msh2Δ* NDT80-AR 1 | 88 | 62 | - | Chr 3 | 2 | 5 | 6 | 1 | 1 |
| *msh2Δ* NDT80-AR 2 | 120 | 108 | - | - | 1 | 4 | 14 | 1 | 2 |
| *msh2Δ* NDT80-AR 3 | 97 | 78 | - | - | 0 | 2 | 6 | 0 | 0 |
| *msh2Δ* NDT80-AR 4 | 81 | 95 | - | - | 0 | 3 | 1 | 0 | 0 |
| *msh2Δ* NDT80-AR 5 | 87 | 82 | - | - | 0 | 1 | 3 | 0 | 0 |
| *msh2Δ* NDT80-AR 6 | 112 | 116 | - | - | 2 | 3 | 17 | 1 | 4 |
| NDT80-AR 1 | 70 | 34 | - | - | 0 | 0 | 4 | 1 | 0 |
| NDT80-AR 2 | 88 | 41 | - | Chr 3 | 1 | 1 | 6 | 1 | 0 |
| NDT80-AR 3 | 113 | 61 | - | Chr 3 | 2 | 3 | 6 | 5 | 0 |
| NDT80-AR 4 | 106 | 62 | - | - | 1 | 2 | 3 | 3 | 0 |
| NDT80-AR 5 | 108 | 43 | - | - | 0 | 2 | 2 | 1 | 0 |
| NDT80-AR 6 | 91 | 51 | - | Chr 5 | 1 | 6 | 2 | 2 | 0 |
| NDT80-AR 7 | 74 | 39 | - | Chr 2, 13 | 1 | 1 | 0 | 0 | 0 |
| NDT80-AR 8 | 106 | 60 | - | - | 1 | 4 | 8 | 3 | 0 |
| *mec1-mn* 1 | 64 | 29 | Chr 2 | Chr 2 | 2 | 4 | 6 | 8 | 0 |
| *mec1-mn* 2 | 96 | 67 | Chr 1 | Chr 1 | 3 | 14 | 13 | 18 | 0 |
| *mec1-mn msh2Δ* 1 * | 66 | 141 | Chr 8 | Chr 8 | 1 | 4 | 7 | 1 | 1 |
| *mec1-mn msh2Δ* 2 | 145 | 195 | - | - | 0 | 16 | 23 | 4 | 6 |
| *mec1-mn msh2Δ* 3 # | 136 | 146 | - | Chr 1 | 0 | 8 | 27 | 1 | 3 |
| *mec1-mn msh2Δ* 4 | 139 | 175 | - | - | 3 | 16 | 23 | 3 | 5 |
| *mec1-mn msh2Δ* 5 | 158 | 153 | - | - | 2 | 18 | 27 | 5 | 4 |
| *mec1-mn msh2Δ* 6 | 148 | 244 | - | - | 3 | 25 | 30 | 5 | 10 |
| *mec1-mn* NDT80-AR 1 | 192 | 125 | - | - | 2 | 9 | 37 | 21 | 0 |
| *mec1-mn* NDT80-AR 2 | 95 | 43 | - | - | 1 | 4 | 6 | 9 | 0 |
| *mec1-mn* NDT80-AR 3 | 131 | 76 | - | - | 2 | 6 | 22 | 15 | 0 |
| *mec1-mn* NDT80-AR 4 | 100 | 73 | - | Chr 14 | 3 | 3 | 13 | 7 | 0 |
| *mec1-mn* NDT80-AR 5 | 147 | 106 | - | - | 2 | 17 | 35 | 32 | 0 |
| *mec1-mn* NDT80-AR 6 * | 63 | 38 | Chr 1 | Chr 3, 4 | 6 | 3 | 4 | 11 | 0 |
| *rad24Δ msh2Δ* 1 | 54 | 46 | Chr 5, 15 | Chr 5 | 0 | 0 | 4 | 0 | 0 |
| *rad24Δ msh2Δ* 2 | 84 | 92 | Chr 1 | Chr 1, 6 | 1 | 9 | 16 | 2 | 4 |
| *rad24Δ msh2Δ* 3 | 104 | 124 | - | - | 0 | 12 | 14 | 2 | 5 |
| *rad24Δ msh2Δ* 4 | 80 | 100 | Chr 6, 8 | - | 1 | 10 | 4 | 0 | 0 |
| *rad24Δ msh2Δ* 5 | 100 | 81 | Chr 8 | - | 1 | 5 | 12 | 0 | 1 |
| *rad24Δ msh2Δ* 6 # | 72 | 97 | - | Chr 3 | 0 | 8 | 6 | 1 | 1 |
| *rad24Δ* NDT80-AR 1 | 140 | 88 | - | - | 3 | 12 | 28 | 17 | 0 |
| *rad24Δ* NDT80-AR 2 | 133 | 54 | - | - | 3 | 7 | 16 | 7 | 0 |
| *rad24Δ* NDT80-AR 3 | 95 | 54 | Chr 6 | Chr 1 3 6 | 6 | 8 | 12 | 7 | 0 |
| *rad24Δ* NDT80-AR 4 | 151 | 104 | - | - | 2 | 13 | 23 | 13 | 0 |
| *rad24Δ* NDT80-AR 5 | 87 | 40 | Chr 6 | Chr 6 | 0 | 6 | 8 | 6 | 0 |
| *rad24Δ* NDT80-AR 6 | 105 | 76 | - | - | 4 | 22 | 20 | 6 | 0 |
| *rad24Δ sml1Δ* | 30 | 11 | Chr 1, 5, 8, 11, 12, 14 | Chr 1, 5, 7, 8, 9, 11, 12, 14, 15 | 2 | 2 | 5 | 8 | 0 |
| *rad24Δ msh2Δ sml1Δ* 1 | 84 | 111 | Chr 3 | - | 0 | 13 | 14 | 2 | 1 |
| *rad24Δ msh2Δ sml1Δ* 2 | 61 | 43 | Chr 3, 8 | Chr 1, 3 | 0 | 6 | 3 | 1 | 0 |
| *sml1Δ* 1 | 67 | 41 | - | Chr 1, 12 | 0 | 1 | 1 | 2 | 0 |
| *sml1Δ* 2 | 82 | 28 | - | Chr 1, 9, 14 | 0 | 1 | 2 | 1 | 0 |
| *zip3Δ* 1 | 65 | 80 | Chr 3 | - | 1 | 7 | 4 | 2 | 0 |
| *zip3Δ* 2 | 46 | 87 | Chr 12 | - | 0 | 6 | 3 | 4 | 0 |
| *zip3Δ* 3 | 55 | 94 | Chr 1 | - | 0 | 5 | 8 | 2 | 0 |
| *zip3Δ* 4 | 44 | 64 | Chr 6 | Chr 5 | 0 | 1 | 9 | 2 | 0 |
| *zip3Δ msh2Δ* 1 | 34 | 306 | Chr 1 | - | 0 | 17 | 10 | 0 | 3 |
| *zip3Δ msh2Δ* 2 | 25 | 295 | Chr 9 16 | - | 0 | 10 | 1 | 1 | 4 |
| *zip3Δ msh2Δ* 3 | 34 | 296 | Chr 15 | - | 0 | 20 | 7 | 0 | 3 |
| *zip3Δ msh2Δ* 4 | 38 | 386 | - | - | 1 | 32 | 10 | 2 | 3 |
| *zip3Δ msh2Δ* 5 | 35 | 382 | Chr 3 4 | - | 0 | 33 | 14 | 2 | 10 |
| *zip3Δ msh2Δ* 6 | 33 | 310 | Chr 11 | - | 0 | 15 | 6 | 0 | 5 |

**Table S2.** Results of separating two component distributions from inter-CO distance distributions. For each genotype, a mixture model was used to extract two component distributions from the distribution of inter-CO event distances. The alpha values approximate the strength of interference in each component distribution, with an alpha of ∼1 being random, and higher values indicating stronger interference. Thus, component 1 has strong interference, and component 2 has a random distribution. The estimated proportion of CO events in each component can be applied to the actual event counts to estimate how many Class I and Class II COs there are in each genotype. The only strain in which two separate distributions could not be extracted was *rad24*Δ *ndt80AR*, which appears to have a single, completely random component. To test the goodness of fit of the two-component distribution to the experimental distribution, a two-sample Kolmogorov-Smirnov test is carried out, with the null hypothesis being that the two datasets derive from the same parental distribution. The comparison for each genotype produces a high P-value of >0.9, indicating that the two distributions there is at least a 90% chance of observing these two distributions if you were sampling a single parent distribution.

| Genotype | $\alpha_1$ | $\alpha_2$ | Estimated Class I % | Estimated Class II % | Mean CO Count | Predicted Mean Class I Count | Predicted Mean Class II Count | P value |
| --- | --- | --- | --- | --- | --- | --- | --- | --- |
| *WT* | 3.04 | 1.09 | 67.5 | 32.6 | 74.7 | 50.4 | 24.3 | 0.9023 |
| *msh2Δ* | 3.81 | 1.19 | 84.3 | 15.7 | 105.0 | 88.5 | 16.5 | 0.9911 |
| *mec1-mn msh2Δ* | 3.56 | 1.37 | 58.7 | 41.4 | 148.5 | 87.1 | 61.4 | 0.9923 |
| *rad24Δ msh2Δ* | 4.25 | 1.34 | 13.5 | 86.5 | 83.4 | 11.3 | 72.1 | 0.9659 |
| *NDT80-AR* | 3.22 | 1.4 | 66.0 | 34.0 | 96.2 | 63.5 | 32.7 | 0.9326 |
| *mec1-mn NDT80-AR* | 3.78 | 1.19 | 44.9 | 55.1 | 137.2 | 61.5 | 75.5 | 0.9635 |
| *rad24Δ NDT80-AR* | - | - | 0 | 100 | 125.0 | 0 | 125.0 | - |

**Table S3.**
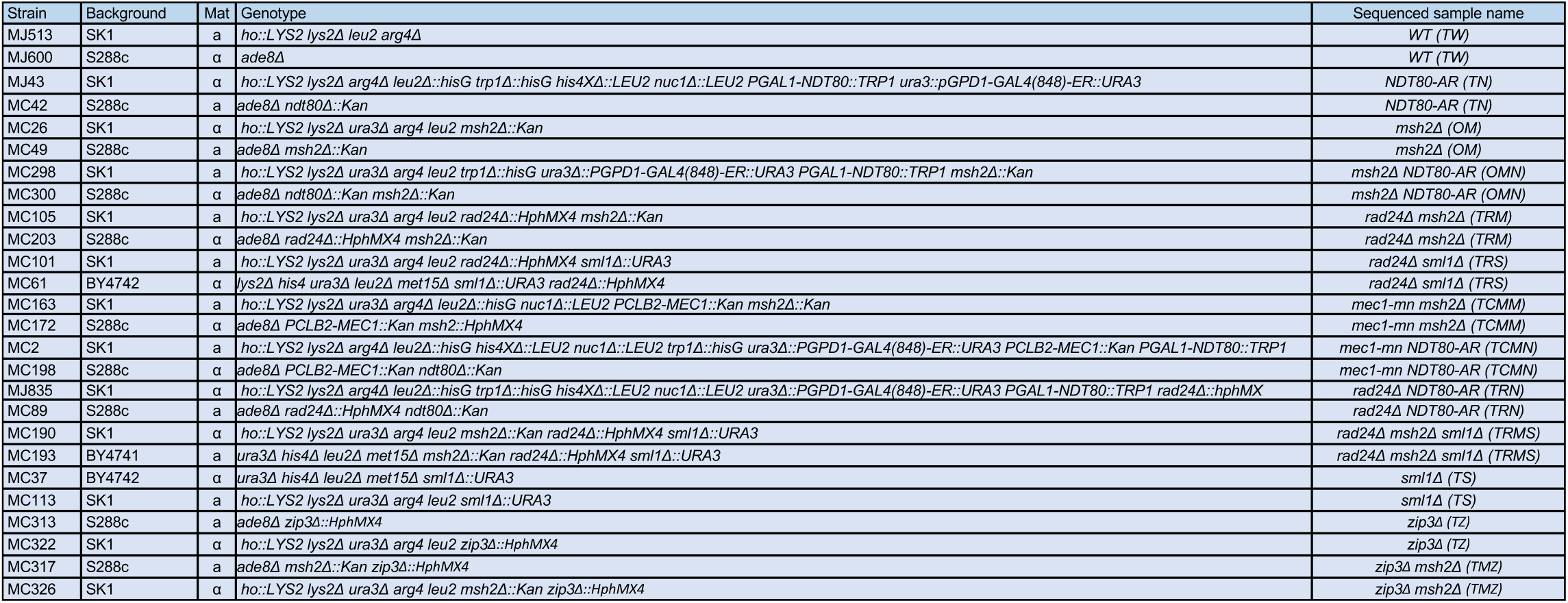
*S. cerevisiae* strains used in this study for tetrad/octad sequencing. All strains displayed are haploid, and were mated immediately prior to sporulation and tetrad dissection.

**Table S4.**
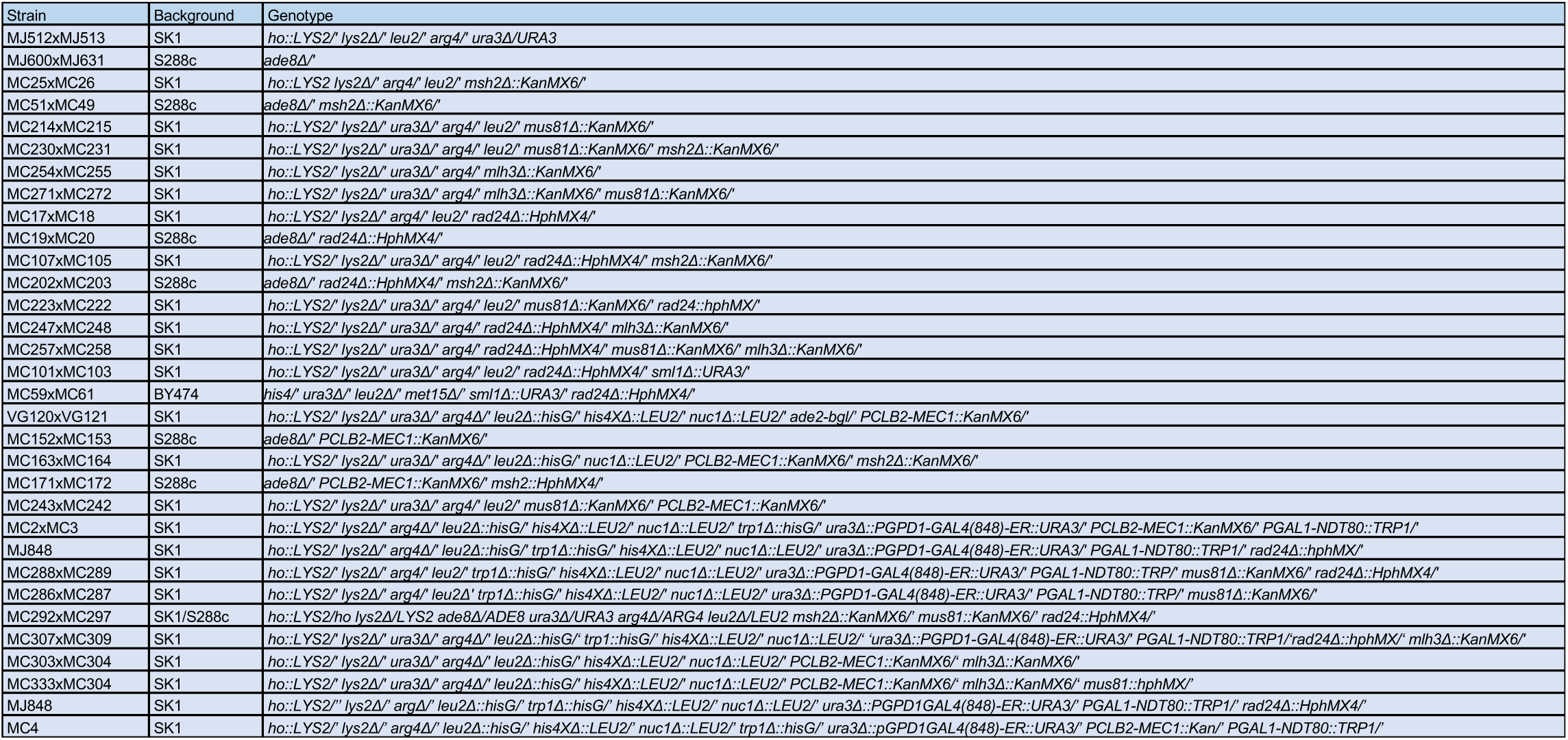
*S. cerevisiae* strains used in this study to measure spore viability. All strains displayed are diploid.

**Table S5.** *S. cerevisiae* spore viability measurements. Viability is scored as the percentage of dissected spores that show visible growth after 48 hours incubation at 30°C. To estimate how well the measurement represents the viability of the population, 95% confidence intervals are used.

| Genotype | Cross | Background | Tetrads dissected | Mean spore viability | 95% confidence interval |
| --- | --- | --- | --- | --- | --- |
| WT | MJ512xMJ513 | SK1 | 64 | 96.9 | 2.1 |
| WT | MJ600xMJ631 | S288c | 72 | 96.9 | 2 |
| WT | MJ512xMJ513 | SK1xS288c | 64 | 82 | 2.7 |
| msh2Δ | MC25xMC26 | SK1 | 110 | 80.9 | 3.7 |
| msh2Δ | MC49xMC51 | S288c | 105 | 77.9 | 4 |
| msh2Δ | MC49xMC26 | SK1xS288c | 149 | 73 | 3.6 |
| NDT80-AR 8h | MJ43xMC42 | SK1xS288c | 23 | 82.6 | 7.8 |
| NDT80-AR 10h | MJ43xMC42 | SK1xS288c | 86 | 70.4 | 4.8 |
| mec1-mn | VG120xVG121 | SK1 | 130 | 58.5 | 4.2 |
| mec1-mn | MC152xMC153 | S288c | 110 | 77.3 | 3.9 |
| mec1-mn | VG120xMC153 | SK1xS288c | 153 | 6.4 | 1.9 |
| mec1-mn msh2Δ | MC163xMC164 | SK1 | 88 | 49.2 | 5.2 |
| mec1-mn msh2Δ | MC171xMC172 | S288c | 111 | 63.1 | 4.5 |
| mec1-mn msh2Δ | MC163xMC172 | SK1xS288c | 176 | 28.8 | 3.4 |
| mec1-mn NDT80-AR 3h | MC4 | SK1 | 66 | 54.6 | 6 |
| mec1-mn NDT80-AR 4h | MC4 | SK1 | 88 | 49.2 | 5.2 |
| mec1-mn NDT80-AR 6h | MC4 | SK1 | 86 | 54.1 | 19.8 |
| mec1-mn NDT80-AR 8h | MC4 | SK1 | 84 | 57.7 | 17.9 |
| mec1-mn NDT80-AR 10h | MC4 | SK1 | 86 | 49 | 12.8 |
| mec1-mn NDT80-AR 4h | MC2xMC198 | SK1xS288c | 64 | 10.2 | 3.7 |
| mec1-mn NDT80-AR 6h | MC2xMC198 | SK1xS288c | 63 | 18.3 | 4.8 |
| mec1-mn NDT80-AR 8h | MC2xMC198 | SK1xS288c | 87 | 16.4 | 3.9 |
| mec1-mn NDT80-AR 10h | MC2xMC198 | SK1xS288c | 63 | 27 | 5.5 |
| rad24Δ | MC17xMC18 | SK1 | 132 | 21.6 | 3.5 |
| rad24Δ | MC19xMC20 | S288c | 106 | 46.7 | 4.8 |
| rad24Δ | MC17xMC20 | SK1xS288c | 240 | 0.7 | 0.5 |
| rad24Δ msh2Δ | MC105xMC107 | SK1 | 107 | 21.3 | 3.9 |
| rad24Δ msh2Δ | MC202xMC203 | S288c | 102 | 28.2 | 4.4 |
| rad24Δ msh2Δ | MC105xMC203 | SK1xS288c | 462 | 5.9 | 1.1 |
| rad24Δ NDT80-AR 4h | MJ835xMC89 | SK1xS288c | 91 | 1.9 | 1.4 |
| rad24Δ NDT80-AR 6h | MJ835xMC89 | SK1xS288c | 93 | 6.7 | 2.5 |
| rad24Δ NDT80-AR 8h | MJ835xMC89 | SK1xS288c | 218 | 7.9 | 1.8 |
| rad24Δ NDT80-AR 10h | MJ835xMC89 | SK1xS288c | 269 | 15.9 | 1.8 |
| rad24Δ NDT80-AR 3h | MJ848 | SK1 | 43 | 22.1 | 6.2 |
| rad24Δ NDT80-AR 4h | MJ848 | SK1 | 44 | 31.8 | 6.9 |
| rad24Δ NDT80-AR 6h | MJ848 | SK1 | 66 | 42.8 | 6 |
| rad24Δ NDT80-AR 8h | MJ848 | SK1 | 84 | 59.2 | 5.4 |
| rad24Δ NDT80-AR 10h | MJ848 | SK1 | 59 | 69.1 | 5.9 |
| rad24Δ sml1Δ | MC101xMC103 | SK1 | 102 | 41.7 | 4.8 |
| rad24Δ sml1Δ | MC101xMC61 | SK1xBY474 | 196 | 4.7 | 1.5 |
| mus81Δ | MC214xMC215 | SK1 | 110 | 55.7 | 4.6 |
| mlh3Δ | MC254xMC255 | SK1 | 110 | 89.1 | 2.9 |
| mus81Δ mlh3Δ | MC271xMC272 | SK1 | 107 | 59.6 | 4.7 |
| mec1-mn mus81Δ | MC243xMC242 | SK1 | 108 | 20.4 | 3.8 |
| mec1-mn mlh3Δ | MC303xMC304 | SK1 | 117 | 49.6 | 4.5 |
| mec1-mn mus81Δ mlh3Δ | MC333xMC334 | SK1 | 94 | 21.3 | 4.1 |
| rad24Δ mus81Δ | MC222xMC223 | SK1 | 110 | 19.3 | 3.7 |
| rad24Δ mus81Δ msh2Δ | MC292xMC297 | SK1XS288c | 31 | 0 | 0 |
| rad24Δ mlh3Δ | MC247xMC248 | SK1 | 109 | 20.6 | 3.8 |
| rad24Δ mus81Δ mlh3Δ | MC257xMC258 | SK1 | 98 | 7.9 | 2.7 |
| rad24Δ mus81Δ NDT80-AR 8h | MC288xMC289 | SK1 | 108 | 9.7 | 2.8 |
| rad24Δ mus81Δ NDT80-AR 4h | MC288xMC289 | SK1 | 89 | 5.9 | 2.4 |
| mus81Δ NDT80-AR 8h | MC286xMC287 | SK1 | 110 | 33.4 | 4.4 |
| mus81Δ NDT80-AR 4h | MC286xMC287 | SK1 | 128 | 34 | 4.1 |
| zip3Δ | MC322xMC313 | SK1XS288c | 106 | 46.2 | 4.8 |
| zip3Δ msh2Δ | MC326xMC317 | SK1XS288c | 177 | 35.9 | 3.5 |

